# A Regression-based Framework for Scalable Pathway-guided Search in Genome-wide Association Studies

**DOI:** 10.1101/241265

**Authors:** Shrayashi Biswas, Soumen Pal, Samsiddhi Bhattacharjee

## Abstract

Traditional unbiased genome-wide association studies (GWAS) have successfully identified thousands of loci associated with various complex diseases but there is evidence to suggest that many variants were missed at stringent genome-wide thresholds. Fortunately, there is a rapidly increasing amount of prior knowledge in publicly available genomic datasets and biological databases that can be harnessed to enhance the power of discovering SNPs/Genes from existing or new GWAS datasets. For most diseases, many of the identified loci tend to cluster into a few specific biological pathways/networks. From the point of view of disease etiology, such clustering is generally to be expected. This phenomenon can be exploited to conduct a more powerful genome-wide scan that is tailored to identify loci that are interconnected in pathways. We propose a scalable regression-based analytical framework to enable such a pathway-guided GWAS and demonstrate that it provides significant gains in power to detect disease associated SNPs. Our method requires two inputs, namely a) genome-wide summary level data (e.g., SNP p-values) and b) a grouping of genes into biologically meaningful categories (e.g., a database of pathways). It automatically adjusts the input p-values by incorporating the knowledge derived adaptively from the data and the pathways specified. The method involves a regularized logistic regression analysis to derive priors of each SNP and then re-weights the p-values of SNPs so as to maximize overall power of making discoveries. It increases the power to discover SNPs co-clustering into some of these pathways, while maintaining the global type-1 error (FWER) at the desired level. We used whole-genome simulations and summary data from real GWA studies of psoriasis, SLE, coronary artery disease and type-2 diabetes to illustrate the power improvement achieved by pathway-guided search. Our pipeline implemented as an R package can flexibly handle large number of prior annotations possibly derived from multiple databases.

## Introduction

Genome-wide association studies (GWAS) have discovered thousands of SNPs (Single Nucleotide Polymorphisms) robustly associated with various complex diseases and traits (1). Although, individually these SNPs have modest effects, collectively they explain a significant percentage of heritability of the diseases/traits (2). Moreover, these SNPs often occur near genes that co-cluster into relevant biological pathways that have helped in elucidating the biology of the disease or trait. Such clustering is generally to be expected if the mechanism of action of the SNPs is through modulation of a nearby gene, as is generally assumed (3,4). Naturally, if this phenomenon of co-clustering of gene-s and hence of nearby SNPs can be assumed a-priori to ‘inform’ the genome-wide search, we expect that power to detect SNPs would improve. More specifically, majority of the SNPs associated with a trait/disease are expected to co-cluster into pathways and hence power to detect such SNPs would improve by borrowing power from each other. For isolated ‘true SNPs’ (less common), power of discovery might be reduced but the overall gain in power should offset this disadvantage. Particularly, for re-analysis of a GWAS data set, such a prioritized search could be very useful in identifying novel SNPs, apart from those already discovered using standard (un-prioritized) GWAS. The novel SNPs thus discovered (if any) would of course need to go through replication and validation steps, similar to the novel findings in the primary scan. Apart from identifying novel loci, such secondary analysis can lend novel insights into the genes and pathways mediating the associations.

Several studies have now shown that trait associated SNPs are enriched in specific functional categories (5–7) such as coding and regulatory regions. Some rigorous statistical methods have been developed for efficient incorporation of ‘enrichments’ of SNP functional annotations to improve power of discovery in GWAS (6,8–11). In the same vein, biological pathways can be thought of as indirect functional annotation categories specified at the level of genes. Pathway enrichment analysis methods are useful for interpreting GWAS hits, but they do not improve the power of discovery. Some authors have considered incorporating pathway knowledge as priors in GWAS of specific diseases using specialized statistical methods or using bioinformatic approaches (12–15). In our opinion, there is still a need for a generic framework that can a) incorporate pathway knowledge in a scalable computationally efficient manner b) is simple to execute and accessible for routine use by the community and c) is modular and extensible so that it can be extended to incorporate different kinds of prior knowledge.

Our objective in this article is two-fold. Primarily, we demonstrate that systematic incorporation of pathway knowledge at the level of genes can indeed improve power to discover associated SNPs in GWAS. Towards this goal, we also introduce an efficient and scalable statistical framework for pathway-guided GWAS and study its properties. Using genome-wide simulations we demonstrate the pathway-guided search does not inflate global false positive rates. Further, using several GWAS datasets as well as simulations we show that considerable power can be gained even without utilizing other knowledge such as SNP functional annotations. In fact, there can be significant power gain simply by agnostically specifying an entire pathway database (i.e. a meaningful grouping of genes) without using knowledge of the biology of the disease to screen for important pathways. It is expected that power can be enhanced further if disease specific knowledge is used carefully.

Roeder and colleagues (16,17) considered statistical methods to incorporate prior knowledge in GWAS from various external sources such as linkage peaks, SNP functional context etc. They pioneered the simple and effective p-value weighting approach which automatically controls global type-1 error (FWER) for arbitrary weights. One difficulty with a naïve weighting approach is the power loss when the assumed prior weights are wrong. They further proposed data-derived adaptive weights that are protected from this kind of prior loss. Similar ideas were discussed in some earlier reports for FDR control (18,19). Since then a number of methods have been proposed for incorporating knowledge of SNP functional annotations based largely on similar principles. Broadly, these methods have two distinct modules, namely ‘Enrichment-Estimation’ of annotation categories followed by ‘Prioritized-Testing’ based on the enrichments. Most existing methods have focused on the ‘Enrichment-Estimation’ module for SNP functional categories by modeling of genome-wide GWAS data or summary results (8,9,20), some have used data or summary results of only the known regions (10,11), e.g., from the GWAS catalog (21). Carbonetto and Stephens (15) proposed a model-based approach for incorporating pathway knowledge but it uses genome-wide individual level data. It also addresses a more complex problem of detecting ‘causal’ SNPs by multivariate modeling. To some extent, methods for prioritizing SNP functional categories also indirectly model causality by explicitly making assumptions such as ‘one causal SNP within a region (8,10,11). In general, these restrictions and causality modeling are appropriate for densely imputed data. In this article, we focus on improving power for discovering ‘associations’ similar to traditional GWA studies, without any joint modeling of SNP-s or sparsity restrictions such as ‘one causal SNP within a region’ (see ‘Discussion’).

For the Prioritized-Testing step, most of the existing methods control FDR or posterior probabilities of SNPs or regions to be true (8,9,15). Recently Steffanson and colleagues (11) proposed different genome-wide thresholds for SNPs based on annotations under global FWER control that is analogous to weighting of p-values (17). Traditionally, in the context of GWAS, FWER (Family-wise Error Rate) has been the accepted standard for type-1 error control. Hence we restrict to the FWER-controlling weighted p-value approach in this study.

Here we build a framework for ‘prioritized GWAS’ using a combination of existing computationally efficient statistical techniques. For the ‘Enrichment-Estimation’ step, our method PMLR (Posterior Marginal Logistic Regression) has an underlying model roughly similar to those of Pickrell (8) or Iversen and colleagues (10) where log-odds of a SNP to be true is modeled as a linear function of annotations. However, being based on (penalized) logistic regression our method is fast and avoids possible issues with instability of convergence that can potentially affect the existing methods requiring full multi-parameter likelihood maximization or MCMC. This can be an advantage of our approach particularly in presence of large number of parameters (i.e. annotations categories). For ‘Prioritized-Testing’ step, we considered simple p-value weights (SPW) proportional to odds of truth, similar to Steffanson (11) and a non-parametric alternative involving optimal cubic p-value weights (CPW), both of which are computationally fast and scalable.

The proposed methods were tested using genome-wide simulations. We observed that global type-1 error is maintained correctly and there is considerable power gain over standard unbiased GWAS in many cases. We also applied them to summary results from 4 GWA studies of Psoriasis, SLE (Systemic Lupus Erythematosus), CAD (Coronary Artery Disease) and T2DM (Type-2 Diabetes Mellitus). In these re-weighted GWAS analyses, we were able to identify multiple ‘crossover’ loci, i.e. loci that were not genome-wide significant in the original GWAS analysis, but became significant after weighting. Further, almost all of these loci have been reported previously from other larger studies, or by combining with more samples in the same study. These were IL13 for psoriasis, 2 loci (near BLK and ITGAM/ITGAX genes) for SLE, at least 2 loci (near PECAM1 and CXCL12 genes) for CAD and 6 loci (near THADA, KCNQ1, ARAP1, MTNR1B, HMGA2 and C2CD4A genes respectively) for T2DM.

In this article, we study the properties of our methods in the context of pathway-knowledge, but the methods are conceptually general and applicable for other kinds of priors such as SNP-level functional annotations or combinations of SNP-level and gene-level priors. Further, these methods should be useful in informing other omics studies such as differential gene-expression or differential methylation studies, with meaningful gene groups (e.g. pathways, ontologies, co-expressed gene sets etc.) as priors. Our pipeline has been implemented in an R package ‘GKnowMTest’ (<u>G</u>enomic <u>Know</u>ledge-guided <u>M</u>ultiple <u>Test</u>ing) that will be made freely available from the R/Bioconductor repository.

## Methods

### Overview of Statistical Methods

#### Model & Notation

Suppose we have summary Z-scores *Z_X_,Z*_2_, … *Z_M_* from a GWAS. Further we have *K* biologically plausible categories of SNPs (henceforth called ‘annotations’). In case of gene-level annotations such as pathways, SNPs are first mapped to genes within a certain base-pair distance (see ‘Bioinformatic Pre-processing’ below) and the genes are then mapped to pathways. This way each SNP is mapped either to a few pathways (through the cis-genes) or remains unmapped. This gives an annotation matrix for the SNPs, *A_jk_* = 1 if the *j^th^* SNP maps to the *k^th^* annotation and *A_jk_* = 0 otherwise. Generally, many SNPs are identical in terms of their ‘map to annotations’ and hence such SNPs are said to belong to an ‘equivalence class’. Let *E*_1_*,E*_2_, …, *E_C_* denote the *C* distinct equivalence classes of SNPs that map to a specific set of annotations. Let *V* denote the *C* × *K* matrix of annotations with only the distinct rows of A (*V_ik_* denotes the binary indicator that the SNPs in the *i^th^* equivalence class all map to the *k^th^* annotation).

We assume that the Z-scores are independently drawn from a 2-groups mixture model, with the ‘proportion of truths’ possibly varying across equivalence classes. Similar to existing methods, we assume an additive logistic model for the annotation effects. Thus, the Z-score of the *j^th^* SNP (belonging to class *E_i_*) is distributed as:

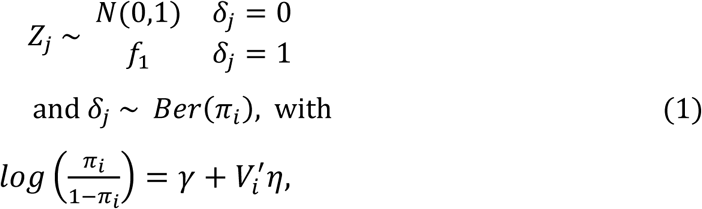
where *γ* denotes the log-odds of a SNP to be associated if it does not belong to any of the annotations and *η* denotes the vector of log-odds ratios contributed by each annotation.

#### Enrichment Estimation by PMLR

We propose to estimate the enrichments (in this case prior probabilities *π_i_*) using a new method PMLR (Posterior Marginal Logistic Regression) involving two sequential steps.

##### Obtaining Posterior Marginals

In the first step, the prior information is ignored and marginal ‘Posterior Probability of Association’ (mPPA) of each SNP is derived. This can be done in various ways such as by fitting a 2-group mixture normal model. Roeder (16,17) assumed *f*_1_ can vary across classes. It is taken to be *N*(*μ_i_*, 1) and *π_i_* and *μ_i_* were estimated using a method-of-moments approach. Bayesian approaches typically assume *μ_i_* = 0 and 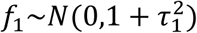 with known specified value of 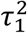 (8,15). We preferred to use a more non-parametric approach following the local FDR method of Efron (22) to avoid restrictive assumptions. Efron’s local FDR works by obtaining the non-parametric density estimate of the marginal distribution 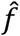 of the Z-scores using a natural-spline or a polynomial to model the histogram bin-counts in a Poisson GLM. The ‘proportion of truths’ is typically estimated by a thresholding or maximum-likelihood approach. The local-fdr for the *j^th^* SNP is given by

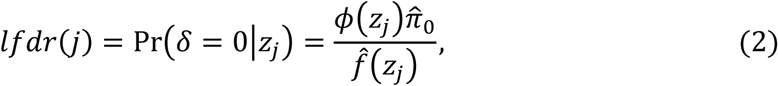
where 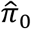 and 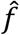 denote respectively the estimated ‘proportion of null SNPs’ and ‘marginal density’ and *ϕ* denotes the standard normal density. We used the R package *locfdr*(23) to derive the *lfdr* values. The mPPA of each SNP is then given by 1 − *lfdr*(*j*).

##### (Penalized) Logistic Regression to derive Priors

To derive SNP prior probabilities, we first obtain average mPPA in each equivalence class 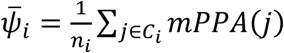 and then use penalized logistic regression of the average mPPA values on the annotations with *n_i_* as weights and a sparseness penalty as follows:

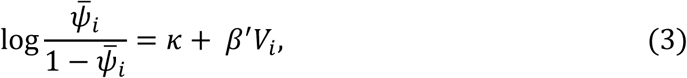
where *V_i_* denotes the *i^th^* row of the annotation matrix V). To fit the parameters we minimize the function below for specific values of *α* and *λ*.

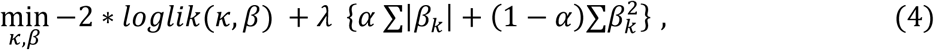
where *loglik* denotes the log-likelihood function based on the logistic model assuming all SNPs to be independent (see ‘Discussion’ section). The above sparse GLM (Generalized Linear Model) can be efficiently fitted using the R package *glmnet* (24) that uses the fast ‘cyclic co-ordinate descent’ algorithm for fitting. It allows the user to control the type of sparseness penalty and degree of shrinkage through the *α* and *λ* parameters. *α* controls the type of penalty (*α* = 1 for LASSO, *α* = 0 for Ridge and intermediate values 0 < *α* < 1 for Elastic NET). *λ* is varied through a sequence of values and the optimal value is either decided by the user or obtained using cross-validation (using the *cv.glmnet* function in R). For the real datasets we used LASSO, Ridge and Elastic-Net returned very similar results (data not shown). Hence we used LASSO (i.e. *α* = 1) throughout for our analyses and simulations. It is possible that the difference across penalization methods may be more relevant when the number of features (i.e. annotations) is much larger than the examples we used. The tuning parameter *λ* (degree of shrinkage/selection for LASSO) is chosen by performing 10-fold Cross Validation using the function *cv.glmnet* and selecting the value of *λ* (from a sequence) with the smallest cross-validation error.

The shrinkage of the *β* values makes sure that even when the number of annotation categories is large or when some categories are small overfitting can be avoided, and the true set of relevant annotations drive the weights. Finally, the shrunk estimates of the SNP prior probabilities are given by the fitted values

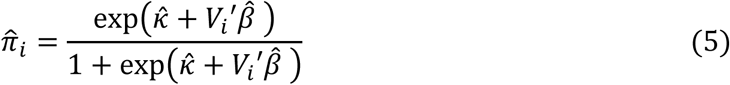

The above procedure is expected to have good theoretical properties under model (**1**) (see **Appendix A**) in estimating the true coefficients (*γ,η*) and prior probabilities *π_i_* of each class.

#### Sequential Priors

In a Bayesian setting, one can propagate priors, in the sense that the posterior from one step becomes the prior for the next step and so on. The PMLR method can be used to incorporate priors sequentially by first running PMLR on the first annotation set using mPPA^(0)^ = 1 Ȓ *lfdr*(*j*), to derive estimated priors 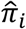 as in Equation (5) and calculating the fitted marginal posterior

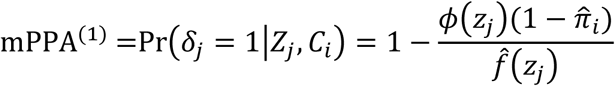

The mPPA^(1)^ is then used for the next PMLR run on the second set of prior annotations and so on. It should be noted that this strategy can give different answers based on the ordering of priors. When possible, it is better to merge all annotations in a single PMLR.

#### Prioritized Multiple Testing

To prioritize SNPs based on prior probabilities of truths 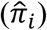 in each annotation group, we use a decision theoretic criterion similar to Roeder and colleagues (17). We consider the maximizing the expected number of true positives among all the tests constraining the expected number of false positives (under the global null) to the desired FWER level (*α* = 0.05). Let *w_i_α* denote the total type-1 error to be allocated for SNPs in the *i^th^* equivalence class. These weights *w_i_* should add to 1 for the FWER to be maintained at *α*. Then the type-1 error spent on each SNP in that class (assuming Bonferroni correction) becomes *w_i_α/n_i_*. Thus a SNP is rejected if *p_j_* < *w_i_α/n_i_*, which is same as saying 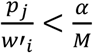, where 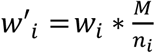 are SNP-level weights with 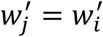 being constant within each equivalence class and satisfying

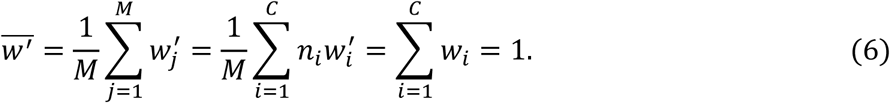

Thus, 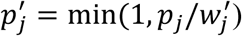 can be considered as a re-weighted p-value, which is the weighted p-value procedure. Roeder and colleagues considered optimizing the average power (expected proportion of true positives), where the normal mean varied in each class. The minor difference in our case is that the alternative CDF *F*_1_ is fixed, but the estimated proportion of truths 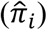 varies across classes. The expected number of true positives is:

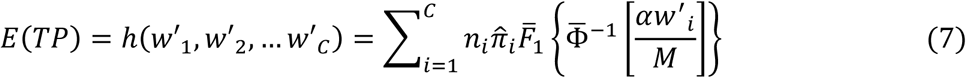

Using a Lagrange multiplier for the type-1 error constraint, we need to maximize.

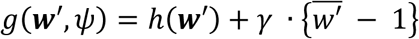
where 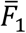 and 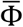 denote the null and alternative upper tail CDFs of the Z-scores. If *F*_1_ is assumed to be *N*(*μ_i_*, 1) or *N*(*μ*, 1), the optimal weights can be easily derived for each fixed *γ* as solution of linear equations (17). We can then solve for *γ* to satisfy the type-1 error constraint. Here we used two non-parametric weighting procedures that do not require the ‘normal with unit variance’ assumption (particularly because we have a fixed alternative distribution unlike the method-of-moments approach of Roeder and colleagues (17)).

##### Cubic P-value Weights (CPW)

Here, we assume the SNP-level weights in a class (at the log-scale) to be a cubic function of the prior log-odds of truth for SNPs in that class, i.e.

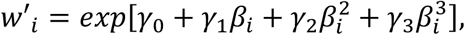
where 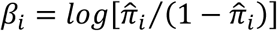. Invoking the constraint that 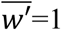, we get

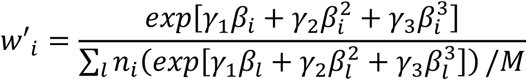

Finally, power (i.e., *E*[*TP*] as in Equation 7) can be maximized for a normal alternative (i.e. *N*(*μ*, 1 + *τ*^2^) or a non-parametrically estimated 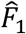 over weights parametrized by (*γ*_1_*,γ*_2_*,γ*_3_), an unconstrained optimization problem. We did limited experiments (data not shown) with higher order polynomials, but did not find significant improvement after cubic exponent.

#### Simple P-value Weights (SPW)

This scheme is essentially CPW with (*γ*_1_ = 1, *γ*_2_ = 0, *γ*_3_ = 0). This is similar to the simple weighting scheme considered previously (11). To see this, we note that odds ratio-s are symmetric (i.e. the prospective odds ratio for a annotated SNP to be causal is equal to the retrospective odds ratio for a causal SNP to map to an annotation. However, an important difference is that, in the current context, odds of a category is based on a multivariate analysis (involving contributions of all pathways). On the other hand, the simple weight described previously is meant for disjoint SNP categories. For a univariate analysis, the methods would coincide.

### Whole-genome Simulations

#### Simulation algorithm

We conducted whole genome simulations using GWASimulator software (25). A crucial modification was required as the software is suited for simulating a small number of causal loci (each located on separate chromosomes). We developed an in-house algorithm for directly simulating a large number of (independent) causal loci retrospectively for a given number of cases and controls. The algorithm (implemented in R) works by simulating a latent variables for causal loci from multivariate normality with different means in cases and controls, and then thresholding the latent variables to get the genotypes (see **Appendix B**). Conditional on the causal locus genotypes, the appropriate phased haplotypes (for the specific chromosome) were sampled (we used Shapeit2 (26) to prephase control individuals of the Psoriasis study). Finally, a modified version of the GWASimulator (C++ code) was used to propagate the genotypes in the flanking regions of the causal SNPs and upto the ends of the chromosomes using a moving window size of 5 SNPs.

#### Type-1 Error & Power

Our simulations were modeled after psoriasis, in the sense that the MAF-s and OR-s of the causal SNPs and genomic locations were chosen to be close to that of 25 suggestive GWAS loci (i.e. p-value < 1e-05) of Psoriasis from the GWAS catalog. The Z-scores of association were calculated in each case using a Cochran-Armitage Trend Test (2-tailed). The simulations were repeated 500 times to check the type-1 errors for various levels by treating SNPs outside ±1 MB of the causal SNPs as null SNPs. This way we had ~500 thousand Z-scores from each of the 500 simulations to estimate type-1 error at small significance levels (effectively somewhat less than 500 thousand, as SNPs close to each other provide correlated information). We used the Z-scores of the causal SNPs (but not neighboring SNPs) to assess power at *α* = 10^−3^ or 10^−5^ or 10^−7^.

#### Pathway Allocation

For the type-1 error and power simulations we used KEGG pathway database. Apart from this we also used synthetic pathways where genes were grouped in a manner so that the degree of connectivity between the causal genes could be controlled (see **Supplementary Methods**).

### Bioinformatic Preprocessing

#### Mapping SNPs to Gene Sets

For any given annotation (e.g. a KEGG pathway), we first mapped the genes to their genomic locations by quering the ‘Transcripts’ table of the hg19 database through the R/Bioconductor package *TxDb.Hsapiens.UCSC.hg19.knownGene* (27). For each transcript we extended it’s range by 500 KB upstream and 100 KB downstream. SNPs were mapped to their genomic locations using the *SNPlocs.Hsapiens.dbSNP142.GRCh37* package (28) containing the SNP identifiers and genomic locations in dbSNP build 142. Finally, the ‘findOverlaps’ function of the *GenomicRanges* package (29) was used to obtain the annotation matrix between pathways and SNPs. Further processing of the ‘findOverlaps’ output was done to obtain equivalence classes for more efficient computation and storage in memory (see **Appendix A**).

#### Deriving KEGG Pathways

We used R/Bioconductor package *org.Hs.eg.db* (30) to generate 229 KEGG pathways (31). The mapping between KEGG pathway identifiers and Entrez gene identifiers was obtained from the ‘org.Hs.egPATH’ object in this package. Entrez gene identifiers were replaced by entrez gene symbols by using the ‘org.Hs.egGENENAME’ object. Names of KEGG pathways was obtained from the *KEGG.db* package (32).

#### Deriving Transfac TF-Target-Gene Sets

We obtained data on transcription factors and their validated targets from Transfac (33). We processed the data from the flat files and used CRAN Package *igraph* (34) to create an igraph object ‘TFG’. Vertices of ‘TFG’ were gene symbols of TF encoding genes and their target genes. An edge in ‘TFG’, say A-> B implies that a TF (with encoding gene ‘A’) targets gene ‘B’. To create Transfac annotations, we considered each TF-encoding gene and all genes in its order 1 neighbourhood as a single pathway. The graph ‘TFG’ had 3228 genes as vertices among which 562 genes are transcription factor encoding genes. We found total 142 different Transfac annotations that had 10 or more genes.

#### Deriving Gene-Ontology (Biological Processes) Annotations

We created GO annotations from GOTERMs of Gene Ontology (35) Biological Process (GO-BP) domain by using two Bioconductor packages *GO.db* (36) and org.Hs.eg.db (30). We used an algorithm to prune the GO-BP tree to create the annotations (see **Supplementary Methods**). The algorithms works by traversing the relatively large (having >50 genes) offspring nodes downwards. The smaller offspring-nodes are not traversed, but merged with their parent node to create an ‘annotation term’. We started with 29586 BP Terms and after pruning we were left with 2570 nodes. Finally, excluding nodes with fewer than 10 or more than 500 genes, we obtained 2326 GO-BP annotations.

#### Deriving a Merged Annotation Set

The gene sets separately obtained from KEGG, Transfac and GO-BP were merged together to give 2697 annotaions. The equivalence classes of each annotation set were used to define a new series of equivalence classes. The advantage of this approach is that the total storage simply adds up (3 separate design matrices), with an extra matrix for mapping between the merged-equivalence class labels to the original equivalence class labels. For example, if the *l^th^* new equivalence class in the merged set denotes SNPs that belong to *i^th^* equivalence class in KEGG, *j^th^* equivalence class in Transfac and *k^th^* equivalence class in GO-BP, then the only additional information stored is *l* → (*i,j k*).

### Results

#### Analysis of Psoriasis GWAS Dataset

We downloaded GWAS data on Psoriasis consisting of 1642 European ancestry subjects (950 psoriasis cases and 692 controls) from the Collaborative Association Study of Psoriasis via the database of genotypes and phenotypes (dbGAP) (37). The original study comprised 1409 cases and 1436 controls. The subset we analysed were those in the ‘General Research Use’ consent group available from dbGAP (dbGaP Study Accession: phs000019.v1.p1). We used Eigenstrat (38) to derive principal components and selected top 4 PC-s to represent ancestry. To run the GWAS, R was used for logistic regression with additively coded SNP genotypes adjusted for principal components.

#### P-value Weights and Shrinkage

Using the summary results obtained from the analysis of Psoriasis data (outlined above), we looked into the behaviour of p-value weights generated by our method with change of penalty (*λ*). For this, we considered KEGG pathways (31) (see ‘**Methods**’) as annotations. The input p-values were transformed to Z-scores as *Z_j_* = Φ^−1^[1 − *P_i_*]. The PMLR method was applied using the R packages *locfdr* to obtain mPPA (Marginal Posterior Probability of Association) values and *glmnet* for penalized logistic regression using LASSO penalty (i.e., *α* = 1). A decreasing sequence of *λ* values was used to derive priors and hence weights (using the Cubic Weighting i. e. CPW method). **Figure 1** shows change in the spread of p-value weights (in the log10 scale) with various *λ* values for LASSO penalty. As expected, the weights have increasing variability around 1 as the degree of shrinkage (*λ*) decreases. For the largest *λ* value, the coefficients are all shrunk to zero and hence the weights are all 1 (i.e. reduces to unweighted analysis).

**Figure 1.**
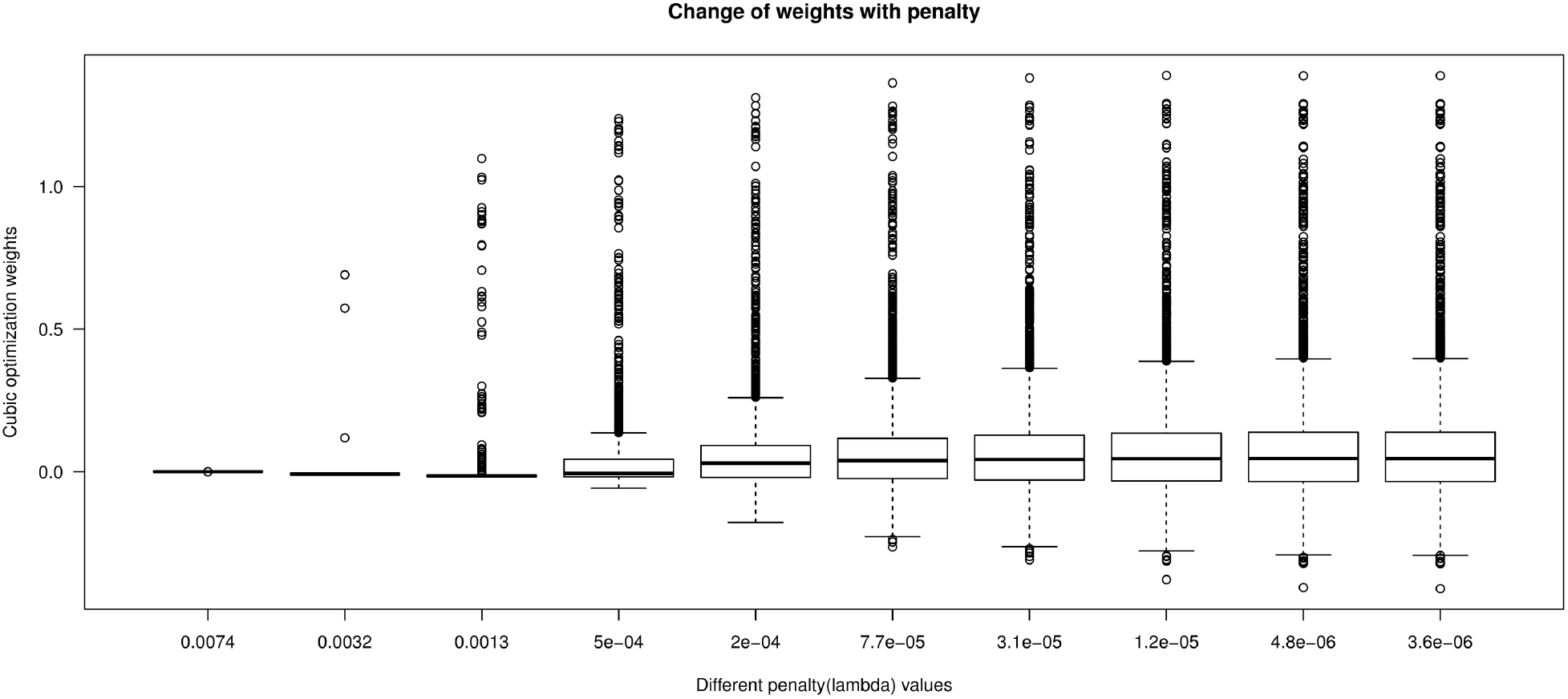
Box plots of weights with change in LASSO penalties. X-axis gives the LASSO penalty λ and Y-axis gives the p-value weights (based on cubic optimization). The plot shows increase in the spread (variability) of p-value weights with decrease in penalty. For the highest penalty all weights are shrunk to 1 (unweighted analysis).

### Simulation Results

In order to study the type-1 error and power properties of our methods we simulated 1500 case and 1500 control genomes based on phased haplotypes having 480493 autosomal SNPs from 690 control individuals in the psoriasis data set (simulation scheme described in **‘Methods’**).

#### Type-1 Error

Firstly, for the 2 weighting schemes namely, Simple weights (SPW) and Cubic weights (CPW) as discussed in ‘Methods’, we checked if the type-1 errors are maintained at different levels starting from 0.1 to stringent levels 1e-07. **Figure 2** gives the bar plot showing type-1 errors of unweighted and weighted p-values for different weighting schemes. It was observed that the type-1 error for both the SPW and CPW schemes are correctly maintained within target levels. It should be noted that 500 replicates are not good enough to estimate type-1 error for any single SNP at low levels of significance, here we looked at only global type-1 error averaged over all null SNPs across the genome (see also ‘**Methods’**). Since the genome-wide average of weights is set to 1 (assuming global null), the average weight among ‘null SNPs’ would generally be below 1 (as true SNPs would tend to get the larger weights) and hence these schemes tend to be slightly conservative.

**Figure 2.**
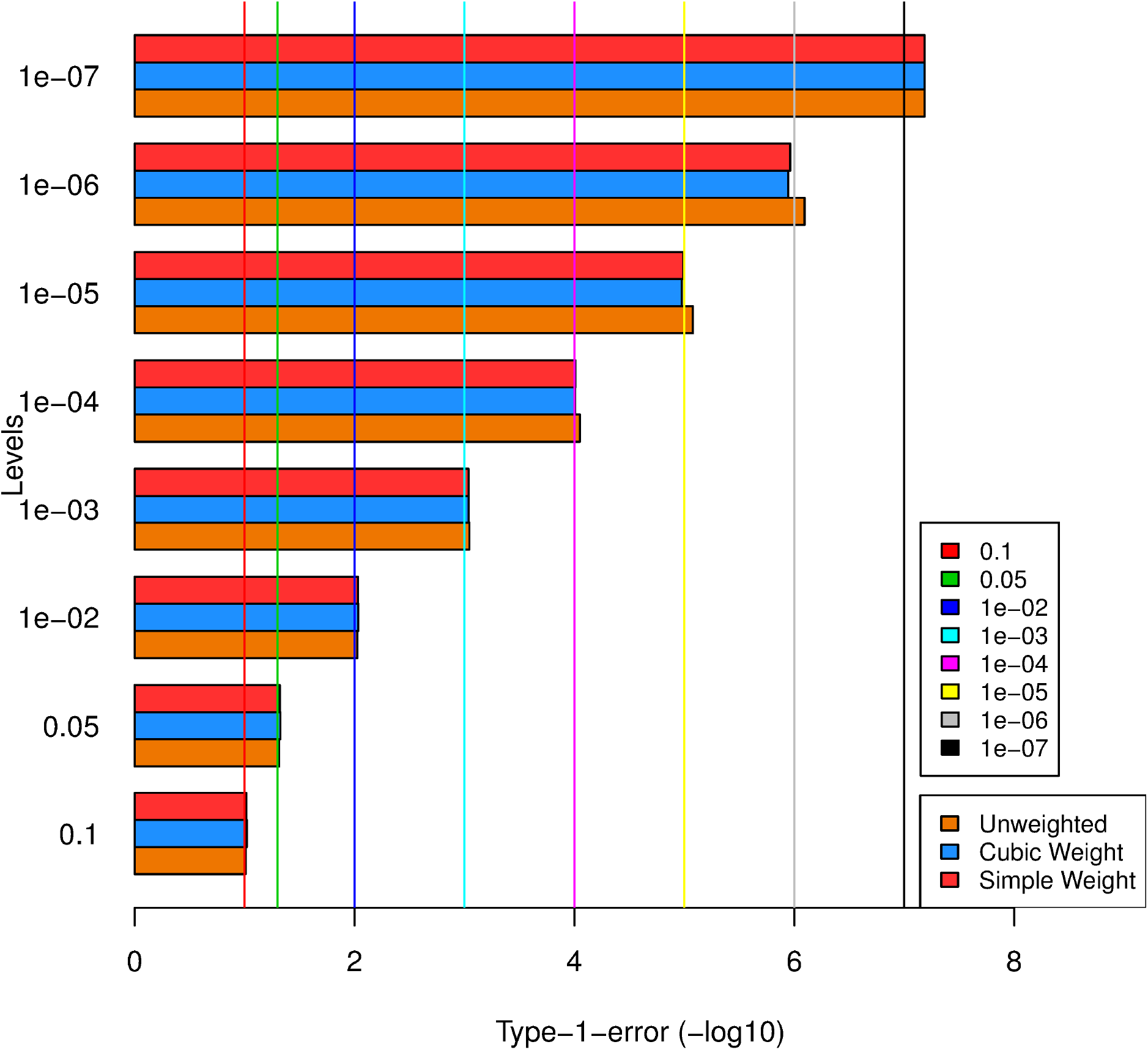
Barplot of type-1 error for unweighted and weighted p-values across levels of significance levels. X-axis shows -log10 values of type-1 error achieved and Y-axis shows the genome-wide significance levels at which each SNPs are rejected (target level). The plot shows that the type-1 error is maintained at different levels of significance.

#### Overall Power

After confirming that the type-1 error is maintained, we looked into the overall power of the weighting schemes using KEGG pathways as annotations. The power was checked for different levels, from 0.1 to stringent GWAS significance level 5e-08. The ‘overall power’ was defined as ‘average number of SNPs crossing the level of significance’ or equivalently the ‘average proportion of simulation replicates in which a causal SNP crossed the level of significance’. Overall power was calculated at different levels for both un-weighted and weighted p-values (SPW and CPW). **Figure 3a** shows the overall power for SPW and CPW at different levels of significance. The difference in overall power between unweighted and weighted analysis increased with level of significance and was highest for level 1e-07. In general, this pattern is expected whenever the sample size is such that most of causal SNP lie below the significance threshold, but not too far from the threshold for weighted analysis to affect its power. Thus, the power improvement at smaller levels (e.g. 1e-04) is small because 1) most of the causal SNPs already have close to 100% power at this level and 2) few of the SNPs have very low power, and hence in spite of up-weighting, they do not cross the threshold. The power of the two alternative weighting schemes is similar, although cubic weights (CPW) performed better than simple weights (SPW) ay lower levels. This is possibly because power of CPW is optimized at a genome-wide Bonferroni threshold.

**Figure 3a.**
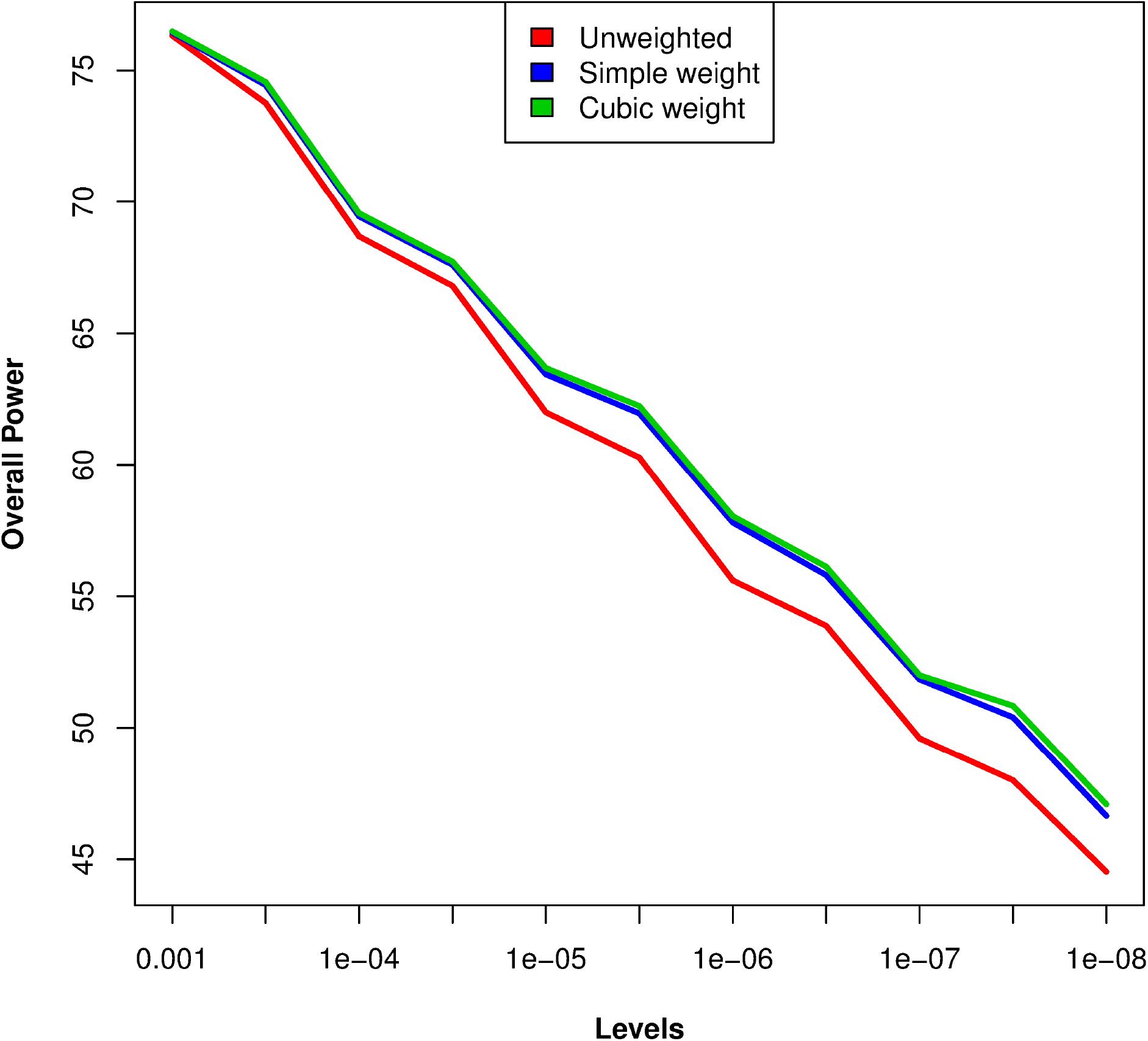
Overall power curve for unweighted and weighted p-values for different weighting schemes at different levels of significance. X-axis shows levels of significance and Y-axis shows overall power (i.e. average power of 25 causal SNPs) based on 500 simulations. The red, blue and green lines show respectively the power for unweighted analysis, simple weighting (SPW) and cubic weighting (CPW).

#### SNP specific power

We also studied the causal SNP specific power from the simulation results. **Figure 3b** shows the bar plot of SNP-wise power for 1) unweighted analysis and 2) weighted analysis (CPW). The powers are shown a genome-wide level of 1e-05 (results for levels 10^−3^ and 10^−7^ were qualitatively similar). In this plot, the causal SNPs are sorted based on decreasing value of difference between power of unweighted p-values and weighted p-values. The plot shows that while some SNPs gain power, most are essentially unaffected while few of them lose power. It is evident from the figure that the gain in power is much more both in terms of magnitude and number of causal SNPs compared to the loss of power. This pattern is to be expected in most realistic scenarios as causal SNPs of a disease would generally map to similar annotations.

**Figure 3b.**
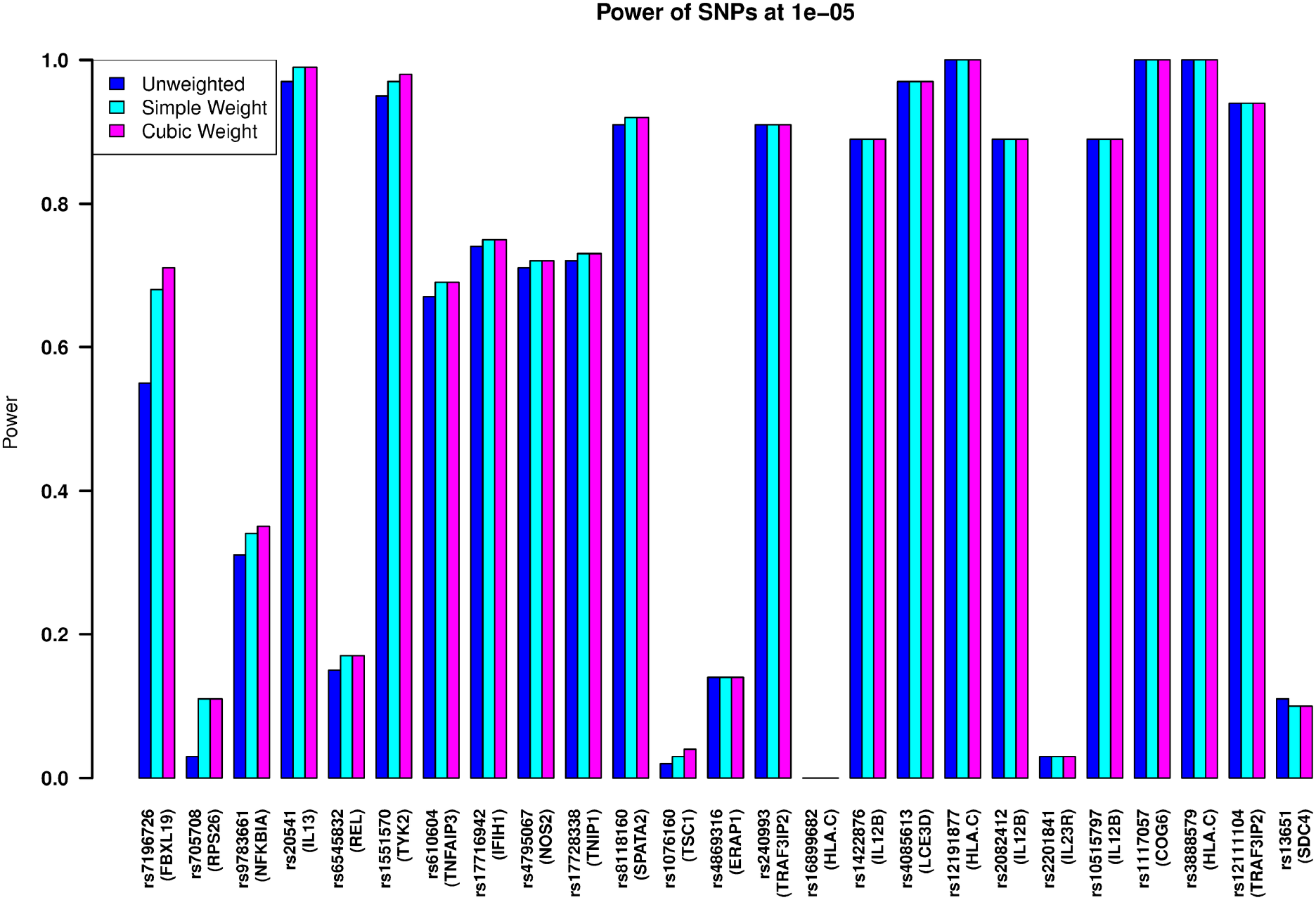
Barplot showing power to detect each causal SNP for different weighting schemes. X-axis shows the 25 causal SNPs and mapping genes and Y-axis shows the power based on 500 simulations. The bars filled with blue, cyan and pink stand for unweighted analysis, simple-weighting (SPW) and cubic weighting (CPW) respectively.

#### Connectedness and Power

It is expected that when a causal gene is ‘more related’ to other causal loci through pathways, it should have higher power to get detected. In order to illustrate that the prioritization method works better when more causal SNPs cluster in few pathways, we generated two example pathway lists as detailed in **Supplemental Methods**. In the first list, we chose *T* (i.e. number of pathways in which the causal genes are distributed) to be 50, so that causal genes are sparsely distributed. On the other hand, the second gene list was chosen with T=10, and hence had more causal genes overlapping within pathways. We found that the overall power of the study increases for the second list, where there is more overlap of causal genes in pathways (T=10). **Figure 4** shows overall power curve across levels for 1) unweighted analysis, 2) KEGG-weighted analysis 3) weighting based on first synthetic pathway list (*T* = 50) and 4) weighting based on second pathway list (*T* = 10). From the figure it is clear that the second pathway list has substantially higher power across all genome-wide levels. In other words, when the causal loci are more connected to each other in pathways, they borrow power from each other and thus the increase the overall power of detecting causal SNPs from weighted analysis.

**Figure 4.**
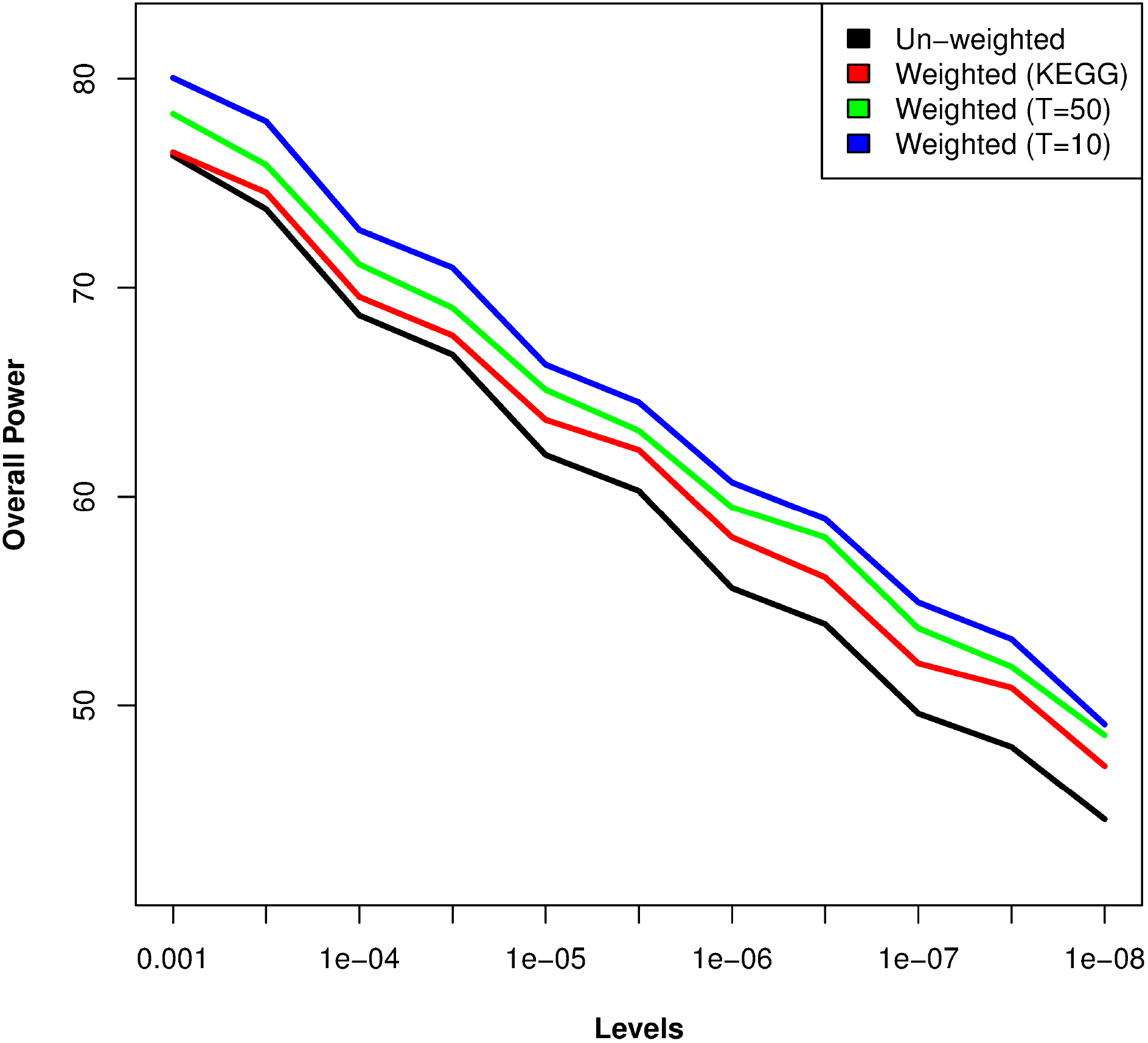
Change of overall power curve with connectivity of genes. X-axis shows levels of significance and Y-axis shows overall power. Black, red, green and blue lines show respectively power of unweighted analysis, KEGG pathways, first synthetic pathway list (T=50, i.e. moderate connectivity) and second synthetic pathway list (T=10 i.e. high connectivity of causal genes).

#### Shrinkage and Power

It is crucial to estimate the degree of penalization from the data. This is because if penalty is over-specified or under-specified it can lead to loss of power. In order to verify this, we estimated power of the causal SNPs for a sequence of *λ* values (the default sequence of *λ* values from *glmnet*) for the LASSO penalty. **Figure 5** shows that the overall power of detection is close to that of ‘unweighted analysis’ for large *λ* values (i.e. low degrees of freedom) and increases gradually with decreasing lambda (i.e. higher degrees of freedom). This is because when *λ* is too large, the weights shrink towards zero, and the PMLR method essentially similar to ‘unweighted analysis’ which has lower power. However, for very small lambda values overfitting results in loss of power particularly at smaller levels of significance. This is expected because when *λ* becomes very small, the PMLR essentially reduces to usual logistic regression without any penalty (shown as ‘glm’) which can unnecessarily put positive weights on ‘small’ annotation categories thereby inflating the variances of the coefficients and lead to reduced power. This effect is prominent at smaller levels of significance. The advantage of regularization is expected to be further pronounced when the number of annotations is very large. The figure shows that weighted analysis (i.e. LASSO with 10-fold CV) avoids overfitting and has the highest power overall across levels.

**Figure 5.**
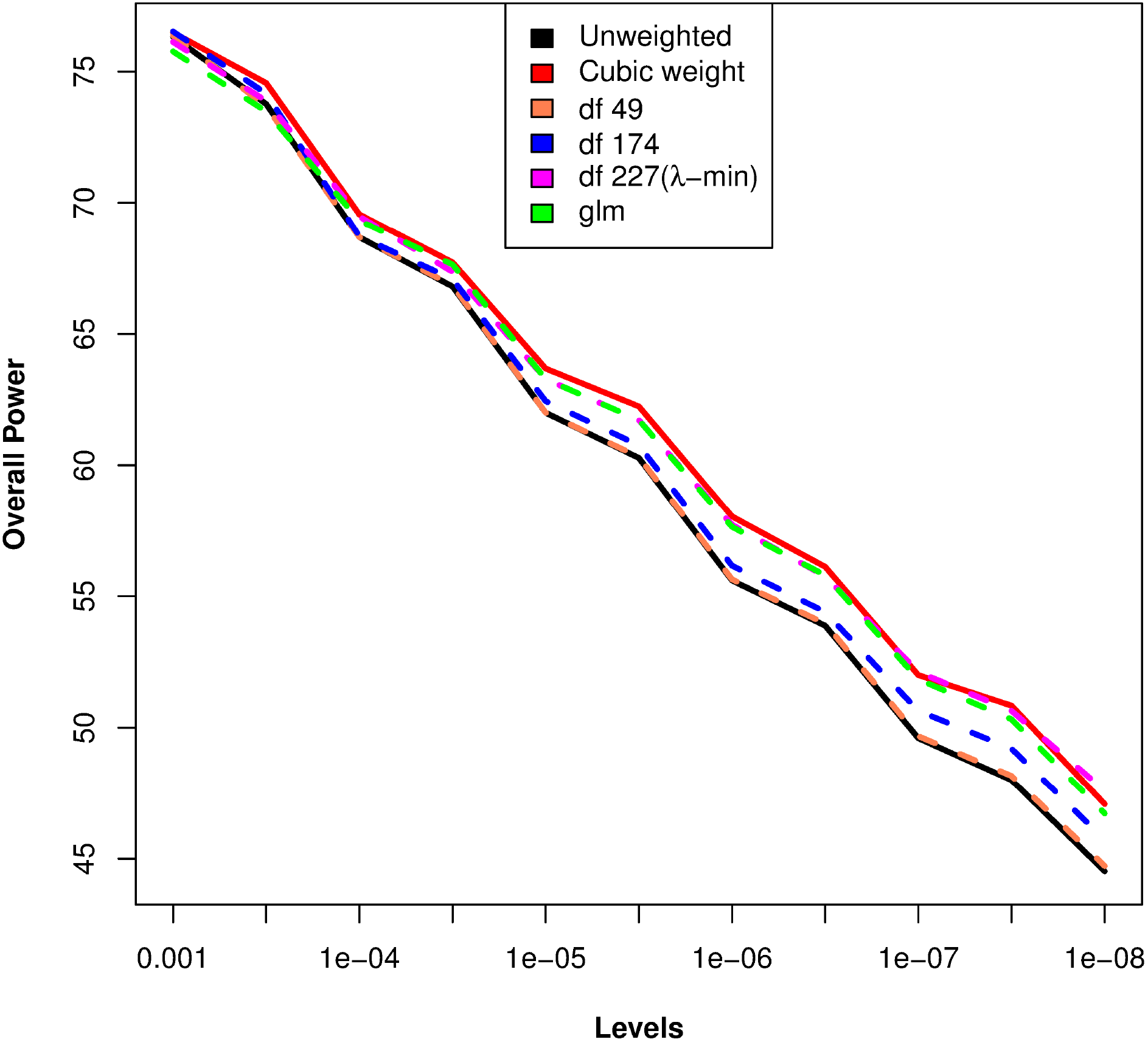
Change of overall power curve with Lasso penalty. X-axis shows levels of significance and Y-axis shows overall power. Black and red solid lines show power of unweighted and weighted-analysis (Lasso with 10-fold CV). The dashed lines show power correspnding to increasing lambda values (decreasing degrees of freedom), namely orange (df=49), blue (df=174) and pink (df=227). The green dashed line shows power for usual logistic regression (‘glm’) without regularization.

### Re-weighted Analyses of GWAS Summary Results

We conducted re-weighted analyses of summary results (i.e. p-values) from four GWA studies using our approach and in each case identified one of more loci (i.e. regions) that were missed by unweighted analysis. For all these datasets, we analysed the Z-scores (derived from p-values as *Z_j_* = Φ^−1^[1 − *P_j_*]) using the ‘Merged’ annotations. We used PMLR with Lasso penalty (10-fold CV) followed by cubic weighting (CPW). Most of these putatively ‘novel’ variants were identified from other studies independently and/or by the same study by using a larger set of samples.

#### GWAS of Psoriasis

We obtained summary results for the psoriasis data on 1677 subjects downloaded from dbGAP (analysis described above). Only 2 loci (HLA-C on chromosome 6 and IL12B on chromosome 5) were found to be significant. Next, we conducted re-weighted analysis of the with KEGG, Transfac, GO (BP) and MERGED annotations separately and inspected 18 known Psoriasis associated SNPs (i.e. p < 5e-08) from the GWAS catalog (21) that were genotyped in our data. **Table 1** shows the unweighted p-values and weighted p-values based on all 4 annotations. All the annotations showed the SNP rs20541 on 5q31 to be genome-wide significant (GWS) i.e., weighted P-values < 5e-08, while the original p-value was 5.98e-07. This SNP did not reach genome-wide significance (GWS) in the discovery phase of the published GWAS which had a considerably larger sample size (comprising 1409 cases and 1436 controls). It was however GWS in the much larger validation stage consisting of 15,369 cases and 19,517 controls (p=5e-09). It has also been found to be GWS in other independent studies (39). **Figure 6a** shows the Manhattan plots of p-values before and after weighting (with ‘Merged’ annotation set). As seen from the Manhattan, in this case the IL13 locus is the only locus that can be considered as a ‘crossover’ locus in the sense that in this region ‘no SNP was GWS’ before weighting while one or more SNPs became GWS after weighting.

**Figure 6a.**
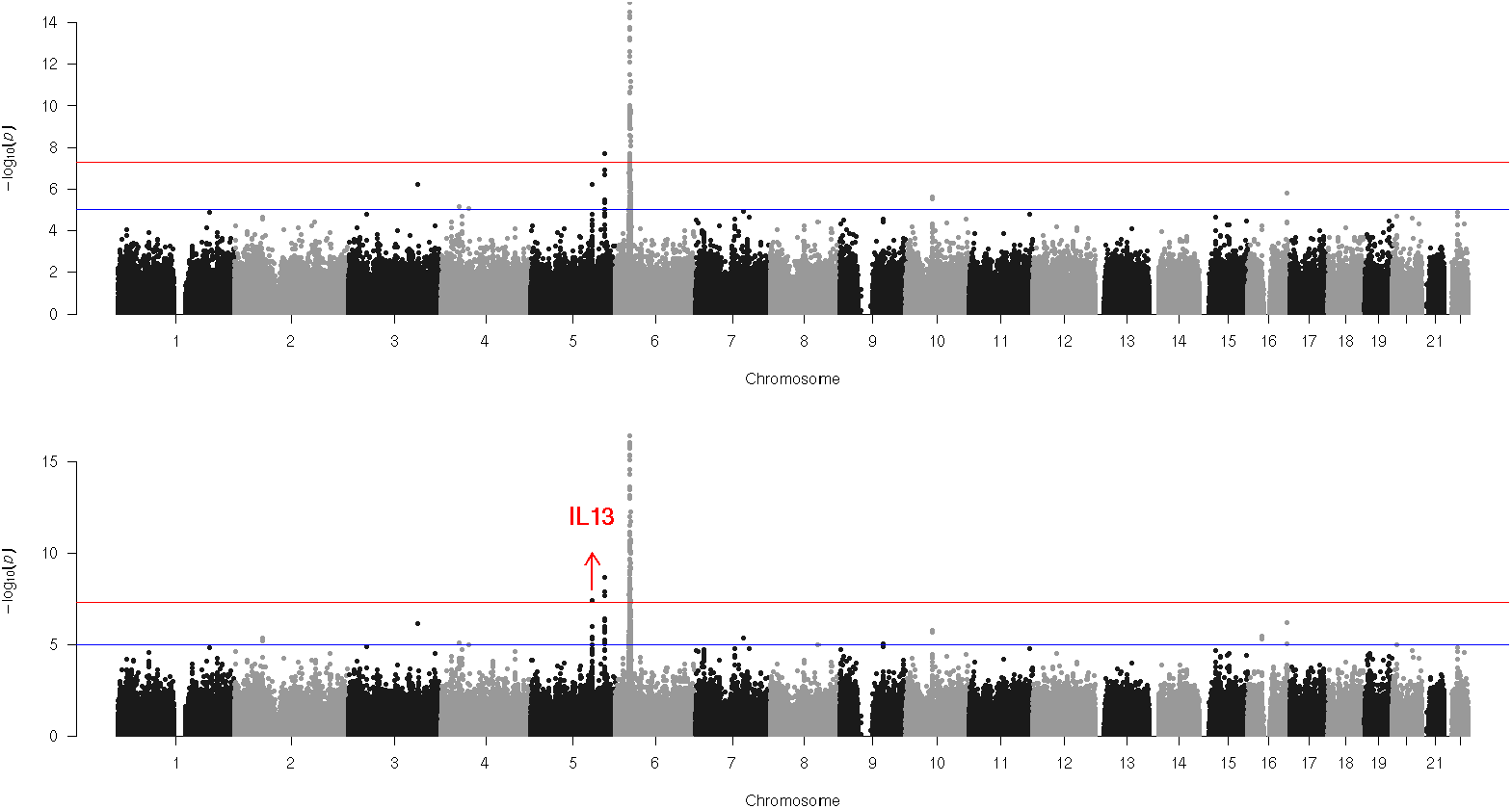
Manhattan plots of Psoriasis GWAS before and after weighted analysis. Upper and lower panels denote Manhattans of unweighted p-values and p-values weighted by ‘Merged’ annotation set (with cubic weighting). IL13 locus is shown as the only ‘crossover’ region (i.e. region newly detected by weighted analysis).

**Table 1.**
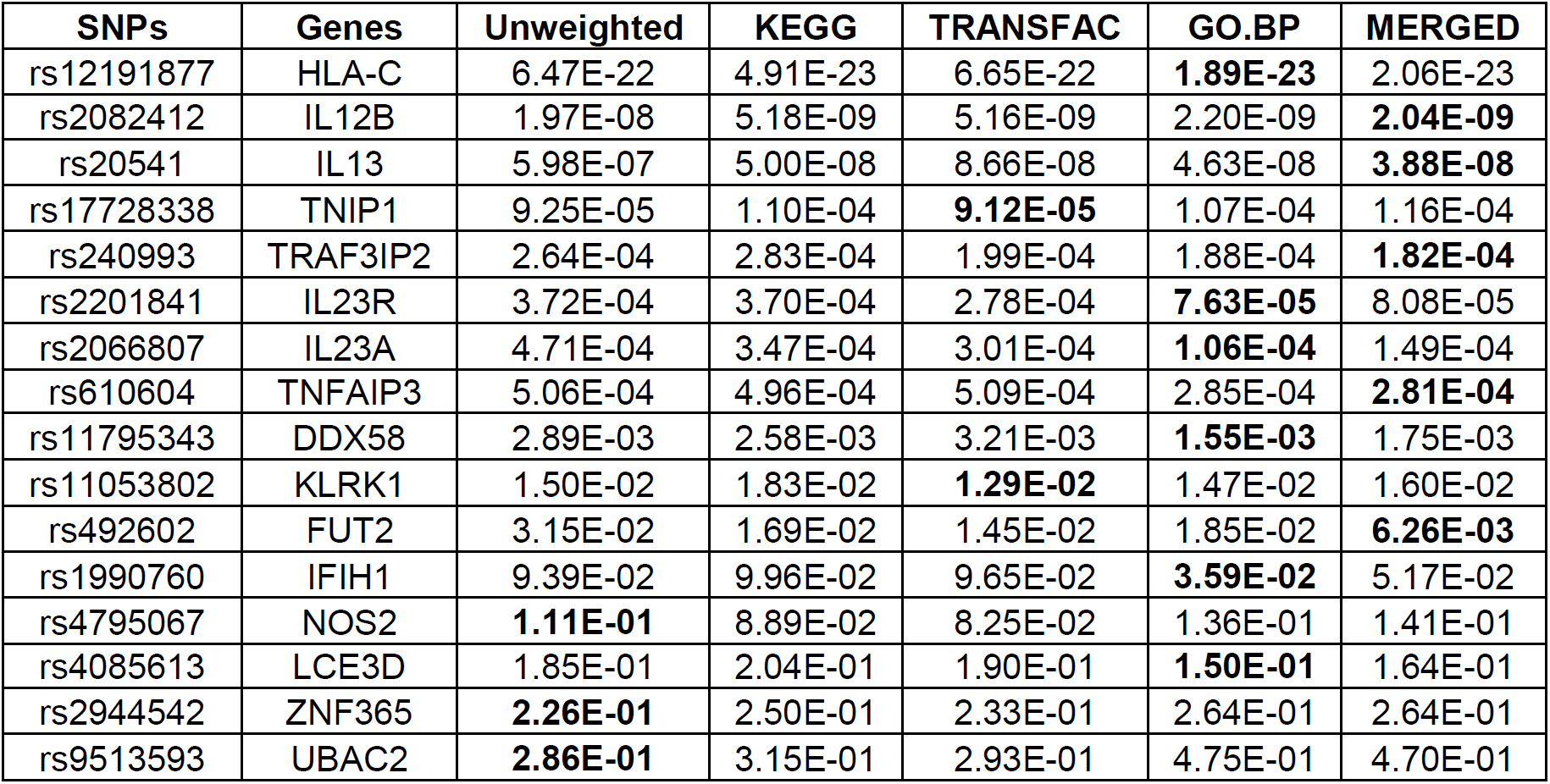
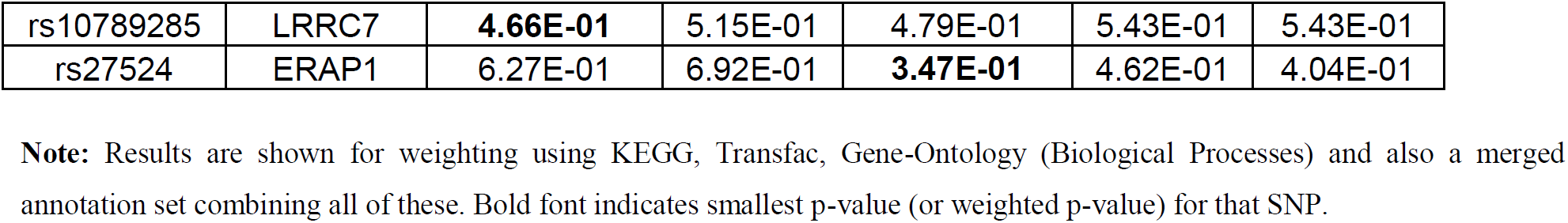
Results of re-weighted analysis of Psoriasis GWAS data for 18 known Psoriasis associated SNPs using 4 different annotation sets.

As evident from **Table 1**, different annotation databases give better p-values for different catalog SNPs in Psoriasis. Overall the ‘Merged’ annotation set gave the best results (smallest (weighted) p-values for 5 out of the 14 SNPs in which p-values improved). This annotation set also avoids biases in favour of previous knowledge about known loci. Hence, we worked with the ‘Merged’ annotations for the remaining weighted-analyses of GWAS datasets.

#### GWAS of SLE (Systemic Lupus Erythematosus)

We downloaded summary data on SLE from the International Consortium on the Genetics of Systemic Lupus Erythematosus (SLEGEN) (40) from dbGAP. These summary data were based on 767 women with SLE and 383 control women available from dbGAP. Unweighted analysis showed 3 GWS loci. Pathway guided GWAS with the ‘Merged’ annotation set gave 2 additional ‘crossover’ GWS loci; one near the BLK gene on chromosome 8p23 and another near the genes ITGAM and ITGAX on chromosome 16p11. The Manhattan plots before and after weighting are showed in **Figure 6b**. The lead SNP in the BLK locus was rs13277113 that has been identified as associated with SLE from other studies (41,42). Similarly, rs4548893 a crossover SNP in the ITGAM locus has been reported previously as GWS (40,41). The unweighted and weighted p-values for these SNPs along with the association results from some independent studies are summarized in **Table 2**.

**Figure 6b.**
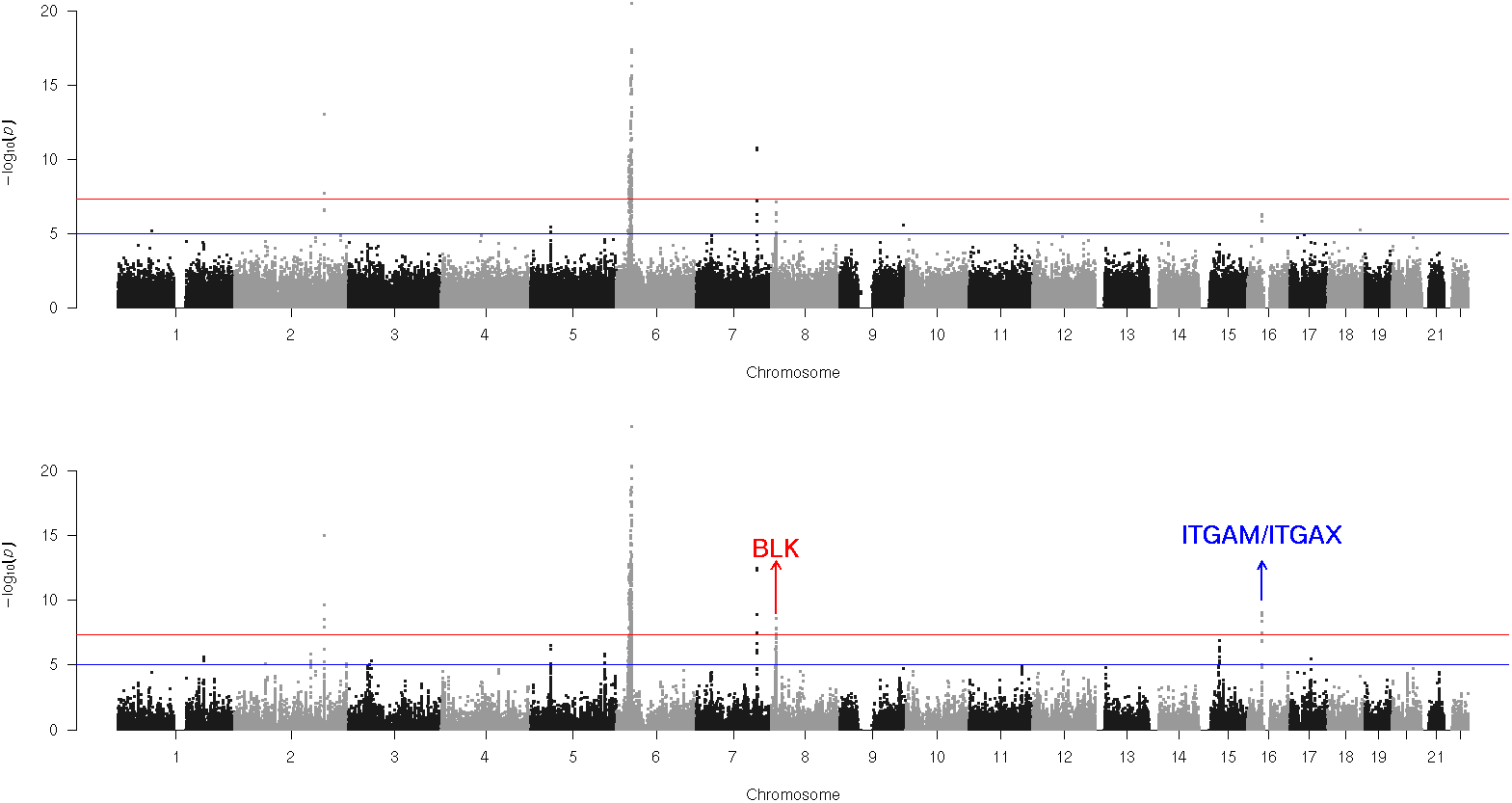
Manhattan plots of Psoriasis SLE (Lupus) before and after weighted analysis. Upper and lower panels denote Manhattans of unweighted p-values and p-values weighted by ‘Merged’ annotation set (with cubic weighting). Two crossover loci (i.e. region newly detected by weighted analysis) are shown with names of genes mapping to those regions.

**Table 2.**
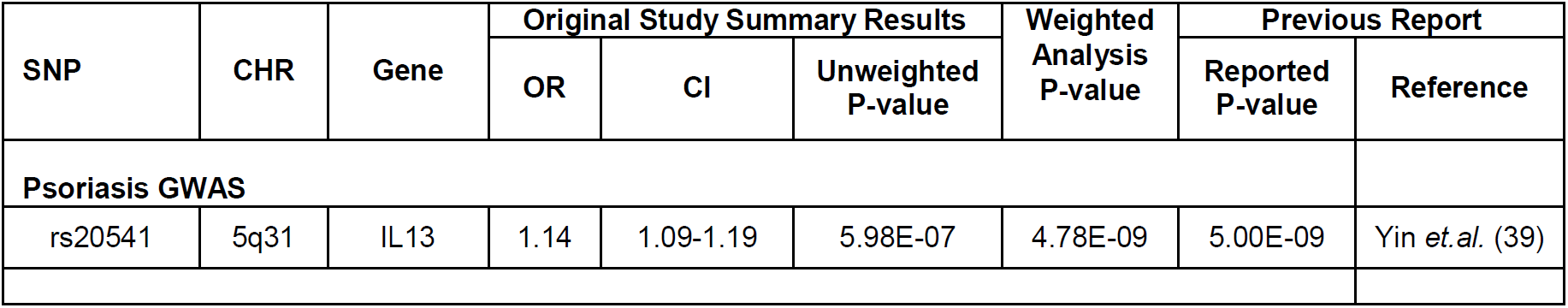
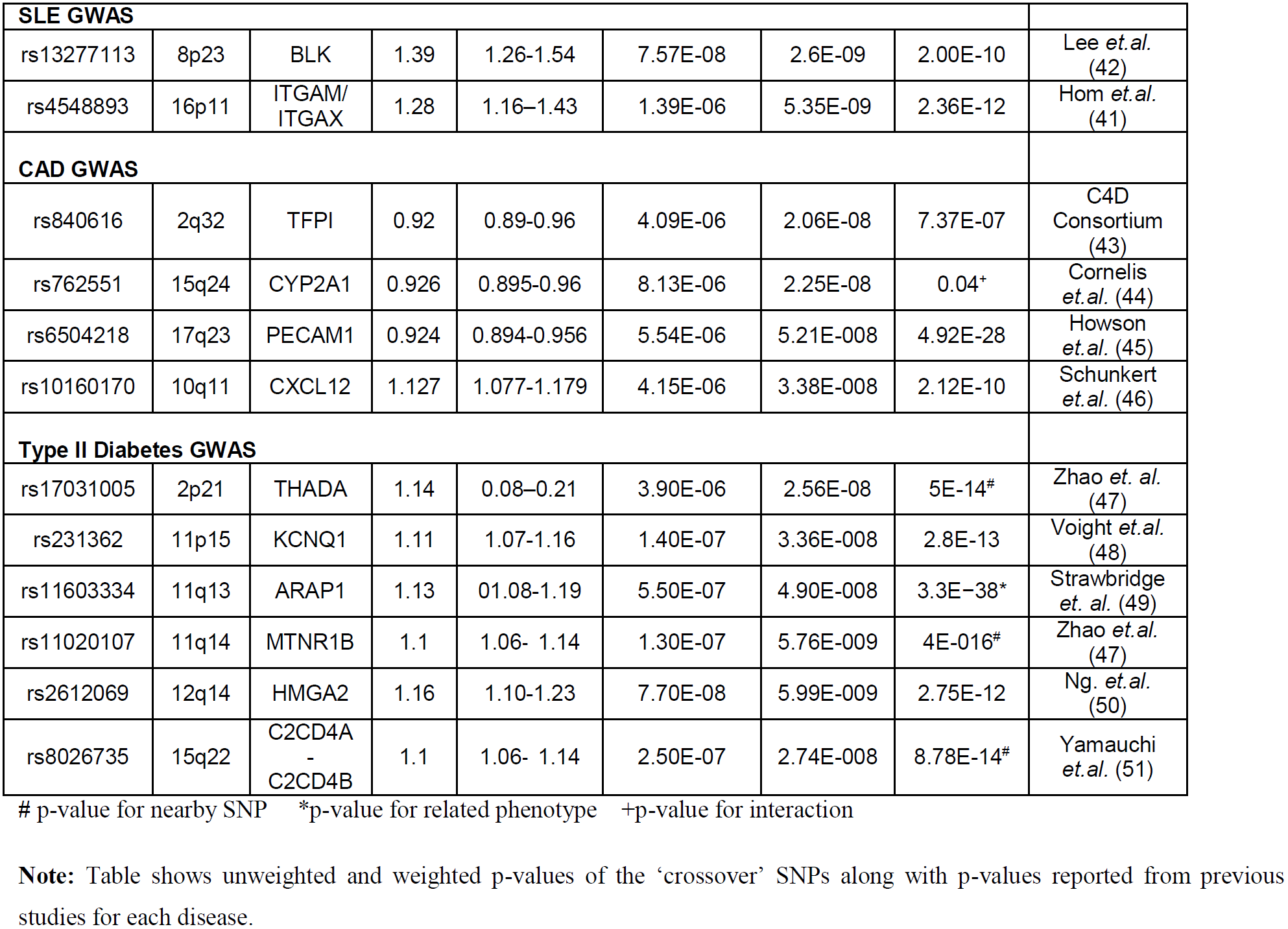
Crossover SNPs from all four GWAS re-analyzed.

#### GWAS of Coronary Artery Disease (CAD)

Summary data on Coronary Artery Disease was downloaded from CARDIoGRAMplusC4D consortium. The data is based on study involving 15,420 cases and 15,062 controls from European and South Asian population (43). Unweighted analysis showed 15 GWA significant loci. Pathway guided GWAS with the ‘Merged’ annotation set gave 4 ‘crossover’ GWAS loci. The Manhattan plots before and after weighting are showed in **Figure 6c**. One crossover locus on 17q23 (lead SNP rs6504218) is near the PECAM1 gene. This region has been identified as a novel locus (lead SNP rs1867624) recently in a larger GWAS meta-analysis comprising 88192 cases and 162544 controls (45). Interestingly these two SNPs (rs6504218 and rs1867624) are in strong LD and the former SNP is an eQTL for PECAM1 in aortic endothelial cells (45). The previously reported SNP rs1867624 was not available in our data. Another crossover locus was on 10q11 near gene CXCL12 (lead SNP rs10160170). SNPs within this region (e.g. rs1746048 less than 300 kb from rs10160170) have been confirmed to be associated with CAD previously (46). The crossover locus on 15q24 is near CYP1A2 gene (lead SNP rs762551). This SNP codes for the CYP1A2*1F allele (164A>C) of the CYP1A2 gene. The ‘C’ allele is known to be a slower metabolizer of caffeine and has potential interaction with coffee intake in CAD (44) and several other phenotypes (e.g., hypertension, Parkinson’s disease, breast cancer etc). However, the marginal association of this allele with CAD (as found here) could be due to residual confounding with other factors such as age, sex and smoking. The fourth crossover locus was near the TFPI gene on 2q32 (lead SNP rs840616). This SNP was reported as suggestive association for the study we analysed (43), but not found in subsequent larger meta-analyses. While it is possible that it is a false positive, it could also have a population specific effect or an interaction with environmental factor(s). Interestingly, the expression of TFPI (Tissue Factor Pathway Inhibitor) has been linked to risk of thrombosis and heart disease previously (52) and a coding SNP in this region (rs7586970 or TFPI N221S within 500 KB) has been shown to be associated with total plasma TFPI levels (53). The unweighted and weighted p-values for the crossover SNPs along with the association results from some independent studies are summarized in **Table 2**.

**Figure 6c.**
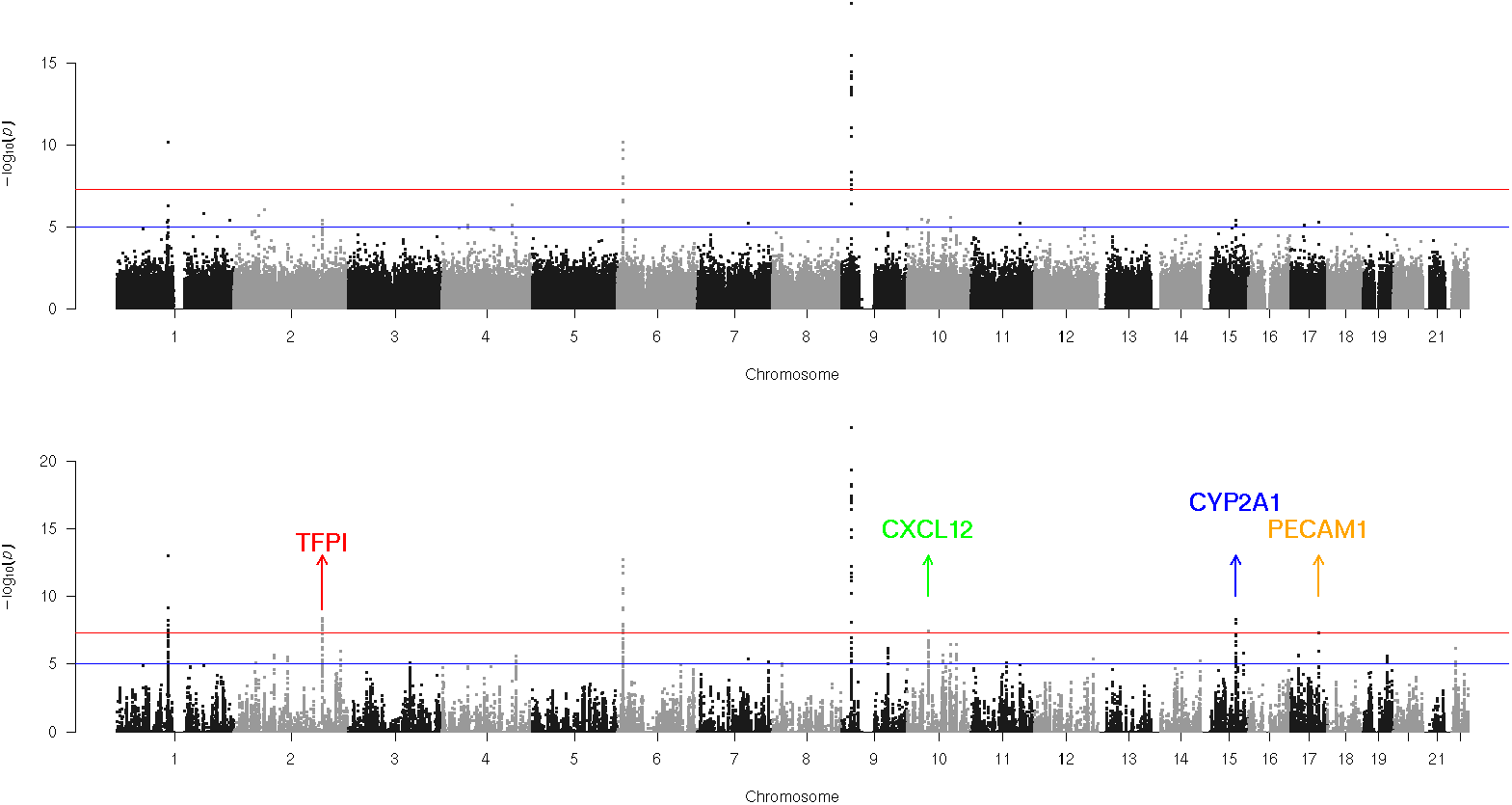
Manhattan plots of Psoriasis Coronary Artery Disease (CAD) before and after weighted analysis. Upper and lower panels denote Manhattans of unweighted p-values and p-values weighted by ‘Merged’ annotation set (with cubic weighting). Two crossover loci (i.e. region newly detected by weighted analysis) are shown with names of genes mapping to those regions.

#### GWAS of Type-2 Diabetes (T2DM)

We downloaded summary data on Type-2 diabetes based from the DIAGRAM (Diabetes) consortium website. These data were based on the stage-1 meta-analysis of 8130 T2DM cases and 38987 T2DM controls of from European population (48). Unweighted analysis showed 12 GWS loci. Pathway guided GWAS with the ‘Merged’ annotation set gave 6 ‘crossover’ loci. The Manhattan plots before and after weighting are showed in **Figure 6d**. One locus near THADA gene on 2p21 had two crossover SNPs (rs10203174 and rs17031005). A SNP in this region rs7578597 (within 100kb of rs17031005) has been previously found to be associated with T2DM (47). Rs10203174 has been reported in a GWAS of ‘platelet count’. Three distinct loci were identified on chromosome 11. The locus on 11p15 was near KCNQ1 gene (lead SNP rs231362). This SNP has been reported in the same article by Voight and colleagues (48) from the Stage-2 GWAS comprising 34,412 cases and 59,925 controls and recently by Zhao and colleagues (47). Another locus on 11q13 was near the ARAP1 gene (lead SNP rs11603334). The SNP has been reported previously as GWS for ‘pro-insulin levels’ (49) and ‘glycated haemoglobin’ (54). The third locus on 11q14 was near the MTNR1B gene (lead SNP rs11020107). A SNP rs10830963 near this gene (less than 10 KB) has been reported previously for association with T2DM (47). A locus on 12q14 (lead SNP rs2612069) was near the HMGA2 gene. SNPs in this region (e.g. rs343092 within 300 KB) have been reported previously for T2DM (50). The last locus was near C2CD4A-C2CD4B genes on 15q22 (lead SNP rs8026735). This SNP is also within 300 KB of previously reported SNP (rs7172432) for T2DM (51). The unweighted and weighted p-values for these SNPs along with the association results from some independent studies are summarized in **Table 2**.

**Figure 6d.**
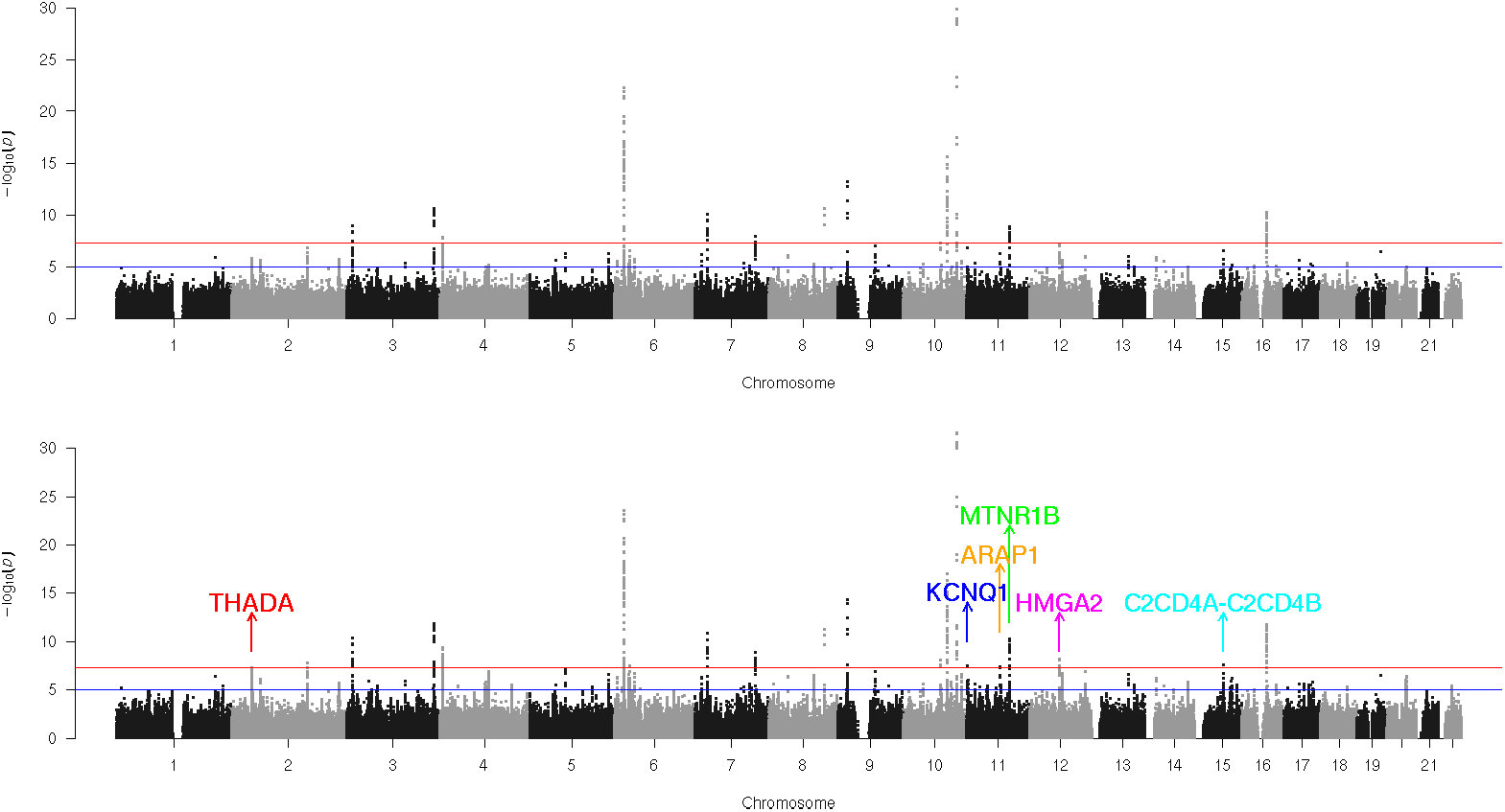
Manhattan plots of Psoriasis Type-2 Diabetes (T2DM) before and after weighted analysis. Upper and lower panels denote Manhattans of unweighted p-values and p-values weighted by ‘Merged’ annotation set (with cubic weighting). Two crossover loci (i.e. region newly detected by weighted analysis) are shown with names of genes mapping to those regions.

#### Significance of Annotations

For each of the above studies we also obtained crude confidence intervals for the KEGG annotations to get an idea about biologically meaningful categories of genes that are enriched for associations with the disease. For each KEGG annotation, we used univariate logistic regression and bootstrap to derive standard errors (see ‘**Supplementary Methods**’). The obtained CI-s for the top 10 KEGG annotations based on p-value are depicted as Forest Plots in **Figure 7**, using the *forest* function of the R package *‘metafor*’. In each case, the top KEGG annotations showing enrichment were largely consistent with common understanding of the etiology of the disease.

**Figure 7.**
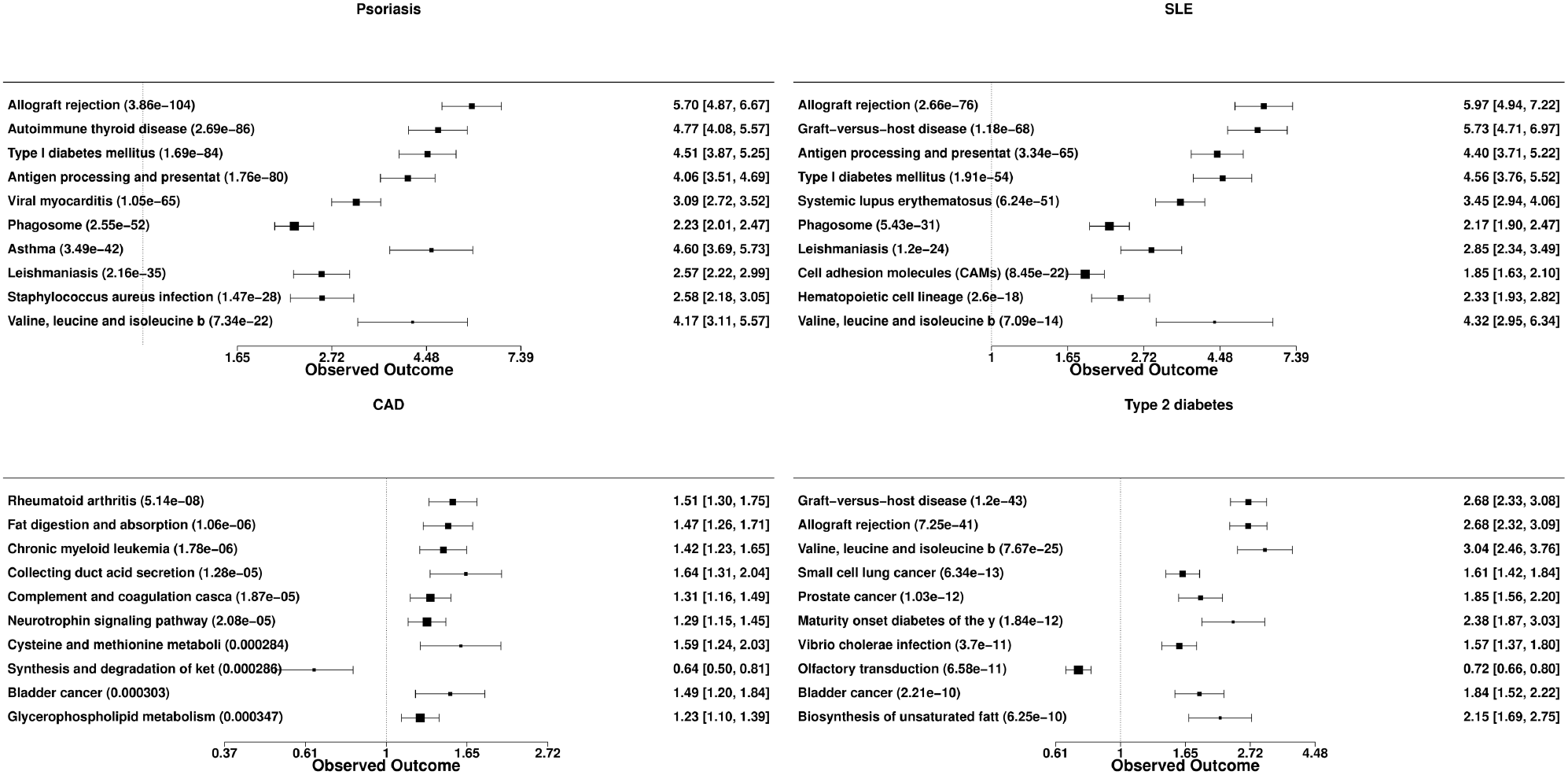
Forest plots showing top 10 KEGG annotations enriched for each disease. The panels shown are a. Psoriasis (top left) b.SLE (top right) c.CAD (bottom left) and d.Type II diabetes (bottom right).

## Discussion

In this article, we illustrate that substantial power improvement can be achieved in GWAS by using knowledge of biologically meaningful categorization of genes, e.g. pathways, ontology terms etc. The power gain is highlighted by the ‘novel loci’ (i.e. crossover loci) detected in the pathway-guided re-analyses of real GWAS datasets. Using, whole-genome simulations, the method is shown to have correct correct global false positive rate for a pathway-guided GWAS, and significant power improvement is observed even with just 25 causal SNPs with realistic effects. For a typical complex disease or trait many more causal variants are expected to be present with more modest effects (55) and this should increase the enrichments and resulting power gain. As expected, there is loss of power for isolated SNPs, but generally such power loss is less common and also much less in magnitude. Moreover, with better prior annotation strategies and as annotation databases become more comprehensive and accurate, these ‘isolated’ true SNPs are likely to be connected with other causal genes, further reducing this concern.

To enable pathway-guided GWAS in a flexible and efficient manner, we discuss a new analysis pipeline. The pipeline consists of a new method called PMLR (Posterior Marginal Logistic Regression) for adaptively estimating ‘enrichments’ (i.e. prior probabilities) followed by optimal p-value weighting to control FWER (Family-Wise Error Rate). The primary advantage of PMLR over existing methods for enrichment estimation in GWAS is that it involves two simple steps, local fdr calculation followed by logistic regression (possibly with a sparseness penalty). Thus it is easily understandable and implementable without requiring significant computational infrastructure or knowledge. It suffices to run a few commands in a statistical package like R. Also, compared to other approaches requiring multi-parameter optimization or MCMC, it is faster, scalable to large number of annotations and also less likely to encounter numerical issues such as non-convergence.

Our method can be interpreted as a single step of an EM-algorithm (as shown in the **Appendix A**). In this sense it can be viewed as making a practical compromise between statistical optimality and computational simplicity. It should be kept in mind that ‘optimality’ only assures good performance under the ‘assumed model’. When optimality of prior estimation under the ‘assumed model’ is desired, the EM can be iterated. However, the final p-value weights depend on the estimated ‘prior probabilities’, and the power depends on the weights, not necessarily on the accuracy of the prior probabilities. Sub-optimal but reasonable ‘prior’ estimates may turn out to be as good as maximum likelihood estimates in terms of eventual weights and power gain.

An obvious and apparent limitation of our approach is that the local FDR approach ignores the correlation structure of the SNPs in deriving the marginal PPA-s. Most existing prior-incorporation methods (e.g. see previous reports (8,10,11,20)) also assume independence across SNPs or SNP blocks. This limitation can be addressed in future by extending our method to allow for correlations among SNPs in the mPPA estimation step (as discussed in **Appendix A**). This highlights the key benefit of a modular apporach. The pipeline can be easily adapted to be used in conjunction with other alternative methods for *mPPA* estimation and/or type-1 error allocation modules.

It should be noted that in this article we consider incorporating gene-level prior knowledge only. Hence, the ‘Enrichment-Estimation’ step does not require assumptions such as ‘one causal SNP’ in a region. This is unlike the Enrichment-Estimation module for incorporating priors on SNP functional categories. This is particularly true as our focus is on discovering ‘associated’ SNPs and secondly, the mapping of SNP-s to genes is done by physical proximity rather than measures of regulatory potential such as those based on pathogenicity (11) or e-QTL associations (56). Hence, in our case, simple *mPPA* calculation ignoring correlations of SNPs and without sparsity restrictions, is adequate for the ‘posterior marginals’ required in PMLR. To combine prior knowledge of SNP functional annotations with pathways, we expect that more sophisticated posterior estimation strategies incorporating ‘local sparsity’ and/or correlations (8,15) (but ignoring priors) followed by PMLR on prior annotations should give a more powerful and optimal approach while retaining the computational efficiency.

In this article, we have refrained from comparing our approach in terms of power to existing statistical methods. The primary reason is the lack of any method, to our knowledge, for incorporating gene-level priors (e.g. pathways) under FWER criterion that is traditionally used for multiple-testing correction in GWAS. Secondly, here our goal was to get an assessment of the power improvement simply from pathway knowledge, un-confounded by power gain from other information such as SNP-level annotations. Most of the SNP-level enrichment methods make assumptions such as ‘one SNP per region’, so using these methods directly for gene-level priors could confound the power gain attributable to pathway knowledge. In future, a careful study of SNP-level annotations and pathways would be required to understand the best way of combining these two sources of information. The logistic regression framework facilitates exploring for non-additive models (e.g., multiplicative interaction) across prior knowledge variables or signatures.

The confidence intervals derived are based on a crude bootstrap procedure that partially mimics the correlation structure of the genome by sampling in blocks. It is possible to give more exact asymptotic variance of the coefficients (from univariate or multivariate model) based on SNP correlation matrix. It is however not straightforward to obtain CI-s for penalized regression particularly when the penalty is estimated from the data without using simulations. This however is not a serious concern. Crude confidence intervals for the regression coefficients (univariate or multivariate) of the selected annotations should usually suffice for the purpose of comparing the relative importance of annotations. There are dedicated methods to obtain enrichment analysis p-values in GWAS that can be utilized if required (57,58).

Here we have considered two p-value weighting schemes to control FWER along the same lines as Roeder and colleagues (17). The weighted p-values although not uniformly distributed under the null hypothesis, can be treated as usual p-values in terms of global (average) type-1 error. We restricted ourselves to type-1 error control, but the PMLR approach can be used to return adjusted FDR estimates. For example, the estimated prior probabilities 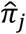 can be used to recalculate posteriors, i.e. local FDR = Pr(*δ_j_* = 0|*z_j_*) or *tail-area* FDR = Pr(*δ_j_* = 0|*Z_j_* ≥ *x_j_*). It is crucial to note that prioritized re-analysis of a GWAS dataset is as likely as the original GWAS to yeild a false positive finding. The chances of false positive may be increased further if an investiagator explores multiple annotation databases untill a discovery is made. To guard against such false-positives, it is recommended that investigators should pre-determine the annotations to be used and also stringent criteria for replication and validation should be followed to confirm a novel finding.

In conclusion, prior knowledge of meaningful ‘gene groups’ such as pathways can be exploited to signficantly improve power of discovery in GWAS. In practice, pathway-guided GWAS can be effectively used for secondary re-weighted scan for additional discoveries. It has potential to synergistically enhance power in conjunction with knowledge of SNP functional annotations. We provide a methodogical framework that can enable routine use of pathway and other annotations for re-weighted analysis of GWAS. Our framework is both statistically rigorous and computationally efficient and scalable. At the same time it is simple and easy to implement using standard GLM and penalized GLM software. It is modular in nature and hence extensible, allowing for future development of further specialized methods for the individual modules.

## Acknowledgements

The authors would like to thank Prof. Partha P. Majumder for valuable suggestions regarding simulations and other experiments and Prof. Probal Chaudhuri for helpful discussion on theoretical aspects. This work was supported by the Wellcome Trust/DBT India-Alliance Intermediate Felloship grant IA/I/14/1/501311 to SB (Samsiddhi Bhattacharjee) and by the National Institute of Biomedical Genomics.

The data/summaries used for the analyses of Psoriasis described in this manuscript were obtained from the database of Genotypes and Phenotypes (dbGaP) found at http://www.ncbi.nlm.nih.gov/gap through dbGaP accession number phs000019. Funding for the Collaborative Association Study of Psoriasis was provided by the National Institutes of Health, the Foundation for the National Institutes of Health, and the National Psoriasis Foundation. Support for genotyping of samples was provided through the Genetic Association Information Network (GAIN). Samples and associated phenotype data for the Collaborative Association Study of Psoriasis were provided by Drs. James T Elder (University of Michigan, Ann Arbor, MI), Gerald G Krueger (University of Utah, Salt Lake City, UT), Anne Bowcock (Washington University, St. Louis, MO) and Gonçalo R Abecasis (University of Michigan, Ann Arbor, MI). For a description of the dataset, phenotypes, genotype data and quality control procedures see Nair et al (2009) Nature Genetics 41:200–204.

The summary results used for the analyses of SLE described in this article were obtained from the database of Genotypes and Phenotypes (dbGaP), at http://www.ncbi.nlm.nih.gov/gap. Genotype and phenotype data for the International Consortium on the Genetics of Systemic Lupus Erythematosus (SLEGEN) (dbGaP accession number phs000216.v1.p1) were provided by Carl D. Langefeld. Funding support for the original study was provided by the Alliance for Lupus Research, the National Institutes of Health, and other sources as detailed in International Consortium for Systemic Lupus Erythematosus Genetics (SLEGEN), Harley JB, Alarcón-Riquelme ME, Criswell LA, Jacob CO, Kimberly RP, Moser KL, Tsao BP, Vyse TJ, Langefeld CD. Genome-wide association scan in women with systemic lupus erythematosus identifies susceptibility variants in ITGAM, PXK, KIAA1542 and other loci. Nat Genet. 2008. 40(2):204–10.

## Appendix A

### Justification of Two-step Prior Estimation (PMLR)

Here we give intuitive justification of the PMLR method only for the case of *K* < *M* and without any regularization. For brevity denote *θ* = (*κ*,*β*) and *X* = [1 *V*]. Assuming *δ*-s to be known, the (complete data) log-likelihood for the logistic model is

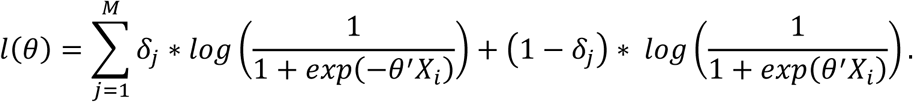

The expected log-likelihood is

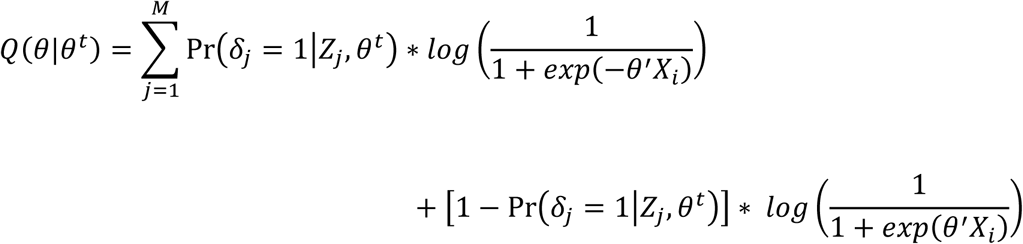

Thus the M-step (without penalty) is simply a logistic regression. In the first step, assuming 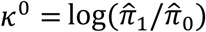 and *β*-s to be 0, i.e. *θ*^0^ = (*κ*^0^, 0, …,0), the Pr(*δ_j_* = 1|*Z_j_*, *θ*^0^) reduces to Pr(*δ_j_* = 1|*Z_j_*) and hence 1^st^ M-step reduces to

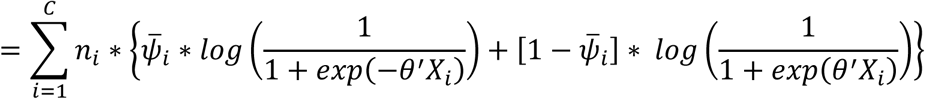

The above expression justifies the reduction of Equation (1) involving SNPs to Equation (3) involving equivalence classes, which gives considerable computational saving (both storage and speed) when *C* ≪ *M*. Both the *glm* and *glmnet* functions allow 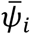 as response with *n_i_* as weights. Clearly the method can be interpreted as 1-step EM starting with a reasonable initial estimate of the posteriors under *θ* = *θ*^0^ (that is itself a solution of an MLE).

Further this posterior becomes more accurate with sample size as the information from the data *Z* dominates that of the prior for large sample sizes. Due to this consistency of the posterior [i.e. 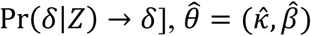 the solution of the logistic regression in Equation (3) converges to the solution of the logistic regression of *δ* on *X*, which is the MLE and hence a consistent and asymptotically efficient estimator of (*γ*, *η*) in **Model (1)**.

### Incorporating SNP correlations in PMLR

Based on the 1-step EM-interpretation above, the PMLR method can be extended to incorporate correlations of SNPs by modifying the *mPPA*-estimation module. In fact, under the full model (with correlations) the complete-data likelihood factorizes as

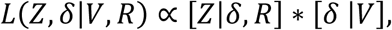
with *V* denoting prior annotations and *R* denoting the correlation matrix for SNP Z-scores. The E-step requires marginal posteriors *mPPA*(*j*) = Pr(*δ_j_* = 1|*Z, R,V*). For the first step of the EM, a Bayesian or Empirical Bayes approach can be adopted to obtain these *mPPA*-s (ignoring prior annotations), i.e. Pr(*δ_j_* = 1|*Z,R*), where *R* is the SNP correlation matrix. This posterior calculation would involve only a few parameters (in absence of prior annotaion variables *V*) and thus the computational efficiency and stability would be maintained. This would be followed by a (sparse) logistic regression as usual. This is because the [*δ_j_*|*V_j_*]-s in the second term are assumed to be independent across SNPs and hence the M-step remains the same (logistic regression of *mPPA*-s on prior annotations *V*). As in the usual PMLR, the method can be iterated for a few steps if required.

## Appendix B

### Retrospective Simulation of Causal SNPs

We assume independent latent Gaussian variables *G_j_*_1_*, G_j_*_2_*, j* = 1, …, *m* corresponding to minor allele indicators for 2 alleles each from ‘m’ causal SNPs, and these are 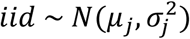 under the additive model. Assume that an individual’s 1*^st^* allele at SNP *j* is the minor allele if *G_j_*_1_ > 0 and absent otherwise. Similarly 2^nd^ allele is minor if *G_j_*_2_ > 0. The parameters *μ_j_* and *σ_j_* are chosen to satisfy the required MAF condition and the equation *E*(*G_j_*_1_|*G_j_*_1_ > 0) − *E*(*G_j_*_1_|*G_j_*_1_ < 0)=1. The model assumed is *logit*[Pr(*D_i_* = 1|*ω_i_*)] = *a* + *ω_i_* where 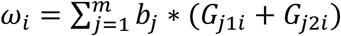. Here *b_j_* is the desired per-allele log-odds ratio of SNP *j* for the usual logistic regression on additively coded genotypes. The intercept *a* is determined by solving for the prevalence of the disease. Note that the population distribution of the latent liability *ω_i_* is normal. First the liability *ω* is retrospectively simulated within cases and controls with its posterior distribution determined by Bayes’ Theorem. Next the latent Gaussian variables *G_j_*_1_ etc are simulated from a (singular) multivariate normal conditional on *ω*. Finally, the simulated Gaussians are thresholded at 0 to give the simulated alleles and genotypes.

## Web Resources

The URLs for the data presented here is as follows.

CARDIoGRAM (coronary artery disease) summary statistics: www.cardiogramplusc4d.org

DIAGRAM (T2D) summary statistics: www.diagram-consortium.org

R package GKnowMTest: https://github.com/sbstatgen/GKnowMTest/

